# Biogeography of Greater Antillean freshwater fishes, with a review of competing hypotheses

**DOI:** 10.1101/2023.09.27.559596

**Authors:** Yibril Massip-Veloso, Christopher W. Hoagstrom, Caleb D. McMahan, Wilfredo A. Matamoros

## Abstract

In biogeography, vicariance and long-distance dispersal are often characterized as competing scenarios. However, they are related concepts, both relying on reciprocal illumination from geological, ecological, and phylogenetic evidence. This can be illustrated for freshwater fishes, which may immigrate to islands when freshwater connections are temporarily present (vicariance), or by unusual means when oceanic gaps are crossed (long-distance dispersal). Marine barriers have a strong filtering effect, limiting immigrants to those most capable of oceanic dispersal. The roles of landspans and oceanic dispersal are being debated for freshwater fishes of the Greater Antilles. We review three active hypotheses (Cretaceous vicariance, GAARlandia, long-distance dispersal) and propose long-distance dispersal to be an appropriate null model due to a lack of reciprocal illumination for landspan hypotheses. Greater Antillean freshwater fishes have six potential source bioregions (defined from faunal similarity): Northern Gulf of México, Western Gulf of México, Maya Terrane, Chortís Block, Eastern Panamá, and Northern South America. Faunas of the Greater Antilles are composed of taxa immigrating from many of these bioregions, but there is strong compositional disharmony between island and mainland fish faunas (>90% of the species are cyprinodontiforms, compared to <10% in Northern Gulf of México and Northern South America, and ≤50% elsewhere), consistent with a hypothesis of long-distance dispersal. Ancestral area analysis indicates there were 16 or 17 immigration events over the last 51 my, 14 or 15 of these by cyprinodontiforms. Published divergence estimates and evidence available for each immigration event suggest they occurred at different times and by different pathways, possibly with rafts of vegetation discharged from rivers or washed to sea during storms. If so, ocean currents likely provide critical pathways for immigration when flowing from one landmass to another, but create dispersal barriers when flowing perpendicular between landmasses. In addition to high salinity tolerance, cyprinodontiforms (collectively) display a variety of adaptations that could enhance their ability to live with rafts (small body size, viviparity, low metabolism, amphibiousness, diapause, self-fertilization). These adaptations likely also helped immigrants establish island populations after arrival and persist long term thereafter. Cichlids may have used a pseudo bridge (Nicaragua Rise) to reach the Greater Antilles, persisting only on the largest islands (Cuba, Hispaniola). Gar (Lepisosteidae) may have crossed the Straits of Florida to Cuba, a relatively short crossing that is not a barrier to gene flow for several cyprinodontiform immigrants. Indeed, widespread distributions of Quaternary migrants (*Cyprinodon*, *Gambusia*, *Kryptolebias*), within the Greater Antilles and among neighbouring bioregions, imply long-distance dispersal is not necessarily inhibitory for well-adapted species, even though it appears to be virtually impossible all other freshwater fishes.

## I. INTRODUCTION

### (1) Intersection of vicariance and island biogeographies

Among biogeographers, mechanisms and processes of how animals populate islands are controversial (Masters et al., 2021; Ali & Hedges 2022). For the Caribbean-Gulf of México region, there is long-standing debate (Ricklefs & Bermingham, 2008; Ali & Hedges, 2021). Three non-exclusive hypotheses are presently studied (Říčan et al., 2013; Roncal et al., 2020) (*i*) Cretaceous vicariance landspan, (*ii*) GAARlandia landspan, and (*iii*) overseas dispersal. Distinguishing among hypotheses should be straightforward. Landspan hypotheses require specific timing and geography because severance of a landspan subdivides formerly contiguous populations on either side of a new barrier. This creates a pattern of vicariance that includes replicate divergence events jointly caused by barrier formation (Platnick & Nelson, 1978; Wiley & Lieberman, 2011). Vicariant faunas that used former landspans should include species incapable of overseas dispersal, in contrast dispersalist faunas that disperse overseas should include only species with exceptional dispersal abilities (Simpson, 1940, 1965). Hence, a pattern of long-distance dispersal, called compositional disharmony, results from dispersal filtering of possible immigrants (Carlquist, 1974). Multi-way filtering occurs where immigrants originate from multiple source areas (Whittaker & Fernández-Palacios, 2007) and this is considered good evidence for dispersal (Platnick & Nelson, 1978). Vicariance biogeography seeks reciprocal illumination between phylogenetic and geologic evidence (Rosen, 1978). A vicariant pattern is supportable when it corresponds in place and time with formation of a dispersal barrier (Platnick & Nelson, 1978; Wiley & Lieberman, 2011). Long-distance dispersal is invoked when divergence occurred across a pre-existing barrier and when cases of dispersal occurred independently by taxon, rather than congruently among taxa (Platnick & Nelson, 1978). Notably, barriers play roles in vicariance and long-distance dispersal. Barrier formation causes vicariance, barrier persistence sets the stage for long-distance dispersal. For this reason, reciprocal illumination applies to both vicariance and dispersal scenarios.

### (2) Freshwater Fishes on Oceanic Islands

As a group, freshwater fishes are very poor dispersers to marine islands. The gap between North America and Vancouver Island appears to be the dispersal limit for this group (Whittaker & Fernández-Palacios, 2007). In this case, one primary freshwater fish (Cypriniformes, *Mylocheilus caurinus*) reached Vancouver Island, likely by virtue of moderate salinity tolerance and facilitated by proglacial flooding from the Fraser River (Clark & McInerney, 1974; McPhail, 2007). Because dispersal along ocean coasts is rare, each drainage discharging to the ocean is a veritable freshwater island (Hugueny, 1989). This dispersal limitation limits species richness in coastal drainages compared to similarly sized drainages that are tributaries to larger rivers (Oberdorff, Hugueny, & Guégan, 1997). For typical freshwater fishes, inter-drainage dispersal occurs via freshwater connections across drainage divides or via stream captures rather than along coastlines or across oceanic gaps (Bianco & Nordlie, 2008; Waters, Burridge, & Craw, 2020). Lands extending from continents to islands provide dispersal routes for freshwater fishes only when mainland and island streams connect. That is, when islands are fully oceanic, with no freshwater connections, only fishes capable of oceanic dispersal can immigrate (Changeux, 1998; Pietsch et al., 2001).

For freshwater fishes, efforts to understand oceanic dispersal have focused on salt tolerance. Myers (1938) and Darlington (1957) classified families largely restricted to freshwater as ‘primary’ freshwater fishes. Families of ‘secondary’ freshwater fishes include taxa often found in brackish water and more tolerant of seawater. Some secondary freshwater families (e.g., Cyprinodontiformes) are adapted to extreme environments while others (e.g., Cichlidae) are more commonly restricted to brackish waters (Bianco & Nordlie, 2008). Families dominated by diadromous fishes are classified separately from freshwater fishes. Comparisons between primary and secondary divisions find that secondary freshwater fishes (as a group) are more broadly dispersed, presumably because they are more prevalent in estuaries (Garcia et al. 2003) and can disperse along coastlines (Matamoros et al., 2015, 2016; Cano-Barbacil, Radinger, & García-Berthou, 2022). However, this classification has been criticized, partly because understanding evolution of salinity tolerance relies on accurate phylogenetic reconstructions (Sparks & Smith, 2005). Progress in this area has revealed that speciation between saltwater and freshwater is common (Seehausen & Wagner, 2014) and can occur repeatedly in secondary freshwater fishes (Ghedotti & Davis, 2013), indicating coastal dispersals can broaden freshwater distributions.

Similarly, biologists working in estuaries recognize fish guilds based on patterns of estuary use (Potter et al., 2015; Whitfield et al., 2022). This includes an estuarine-freshwater guild (species that form populations in either habitat) and a freshwater estuarine-opportunist guild (species that populate estuaries under fortuitous conditions). Freshwater fishes that sporadically occur in estuaries, usually in limited numbers, are freshwater stragglers. These guilds further illustrate that only a subset of freshwater fishes can maintain populations in coastal habitats, including secondary freshwater fishes like Cichlidae and Cyprinodontiformes (Whitfield, 2015).

Because families classified as primary freshwater fishes include some species with substantial salt tolerance and marine distribution (Sparks & Smith, 2005), salt tolerance may not entirely explain why secondary freshwater fishes are more prevalent on islands. Based on a wide range of organisms (not including fishes), Thiel & Gutow (2005) noted that organisms rafting through the ocean are often asexual or have internal fertilization, with internal incubation of embryos or deposition of eggs on vegetation, and small body size. Accordingly, within Cyprinodontiformes, selfing representatives of Rivulidae (a family of secondary freshwater fishes) are widespread throughout the Antilles (Tatarenkov et al, 2017), as are live-bearing representatives of Poeciliidae (another family of secondary freshwater fishes; Reznick et al., 2017).

Excellent examples of immigration by primary freshwater fishes via continent-island drainage connections are documented on the eastern coasts of Asia, where there are extensive continental shelves that allow for river-drainage connections between the continent and islands during sea-level falls (Voris, 2000). For instance, streams draining the islands of Hainan and Taiwan integrated with mainland drainages during Pleistocene sea-level low stands and freshwater fishes populating both islands were a random subset of fishes available from mainland source pools (Lin et al., 2023). This included primary freshwater fishes, notably Cypriniformes (Chiang et al., 2013; Ju et al., 2018; Wang et al., 2022; Chen & Jang-Liaw, 2023). Similarly, Late Miocene-Early Pliocene drainage connections allowed cypriniform fishes to populate the Japanese archipelago (Tominaga, Nakajima, & Watanabe, 2016; Jang-Liaw et al., 2019; Taniguchi et al., 2021).

Elsewhere, lowered sea levels resulted in drainage integration between the Indian Peninsula and Sri Lanka (Ramasamy & Saravanavel, 2019), allowing cypriniform fishes to populate (Sudasinghe et al. 2020, 2021a, 2021b). During the Last Glacial Maximum, the Rhine River extended through the exposed English Channel, integrating streams of unglaciated Great Britain (Hijma et al., 2012). Cypriniform fishes populated these streams, but not unconnected ones farther north and west (Wheeler, 1977). Lower sea levels in the Ionian Sea allowed integration of tributaries to the Gulf of Patras (Perissoratis & Conispoliatis, 2003), which facilitated immigration of cypriniform fishes to western rivers on the Peloponnese Peninsula (Durand et al., 1999; Buj et al., 2019). Lowered levels in the Adriatic Sea allowed seaward expansion of the Po River, integrating drainages isolated at higher sea levels (Monegato et al., 2015). This enabled cypriniform fishes to disperse broadly (Ketmaier et al., 2004) and populate Pag Island (Vucić et al., 2023). These examples demonstrate reciprocal illumination. Geological evidence supports the existence of freshwater connections that could have facilitated dispersal of freshwater fishes to future islands and peninsulas, and presence of primary freshwater fishes on formerly connected islands and peninsulas is congruent with this geological evidence.

### (3) Freshwater Fishes in the Greater Antilles

Myers (1938) noted that all native freshwater fishes in the Greater Antilles are secondary freshwater fishes, mostly within Cyprinodontiformes (Poeciliidae, Rivulidae, Cubanichthyidae, Cyprinodontidae, Fundulidae). The only exceptions to this pattern are one lineage each from Cichlidae and Lepisosteidae (Table 1). As a group, Cyprinodontiformes includes many taxa with unique adaptations to withstand harsh environments (*i*) amphibiousness (Turko & Wright, 2015), (*ii*) tolerance of elevated hydrogen sulphide (Greenway et al., 2020), (*iii*) viviparity, placentotrophy, and superfetation (Pollux et al., 2009; Helmstetter et al., 2016), and (*iv*) self-fertilization (Avise & Tatarenkov, 2015). Numerous species exhibit high tolerance for extremes and fluctuations in temperature, dissolved oxygen, and salinity (Nordlie, 2006). Cyprinodontiforms also have low metabolic rates (Nordlie, 2014) and can lower metabolism in stressful conditions (e.g., *Limia melanonotata* Poeciliidae of Hispaniola, Haney & Walsh, 2003). Island populations of cyprinodontiforms appear to have slower metabolisms than continental relatives (Nordlie, 2014), possibly because a lower metabolism was an aid in oceanic dispersal. Thus, inherent capacities to tolerate harsh, fluctuating environments and adapt to environmental conditions that are marginal for other freshwater fishes could account for the unrivalled success of Cyprinodontiformes in populating the Greater Antilles.

**Table 1.** Summary of best available information for freshwater fish immigrations to the Greater Antilles from adjacent bioregions (ancestral areas). Taxa are listed in tentative chronological order for timing of immigration, based on available divergence estimates (origin, references provided in text). Ancestral areas: Northern South America (NSM), Maya terrane (MT), Northern Gulf of México (NGM), Chortís block (CB), Western Gulf of México (WGM), Eastern Panamá (EP). Underlined areas were not resolved in the ancestral-areas analysis, but are indicated in evidence from literature review. Distribution of taxa among islands of the Greater Antilles is given. Alternative scenarios for the *Limia*-Hispaniolan *Poecilia* clade are provided (shaded rows). Hisp. = Hispaniola, Jam. = Jamaica, PR = Puerto Rico, CI = Cayman Islands, BI = Bahamas Islands.

| Taxon | Origin (Ma) | Dispersal or vicariance | Ancestral areas | Cuba | Hisp. | Jam. | PR | CI | BI |
| --- | --- | --- | --- | --- | --- | --- | --- | --- | --- |
| **Paleogene (66.0-23.0 Ma)** |  |  |  |  |  |  |  |  |  |
| *Quintana-Girardinus* (Poeciliidae) | 50.7-22.3 | Dispersal | NSM, EP, MT, WGM | X |  |  |  |  |  |
| *Nandopsis* (Cichlidae) | 29.4-23.2 | Either | NSM, CB | X | X |  |  |  |  |
| *Limia* + Hispaniolan *Poecilia* (Poeciliidae) – SINGLE IMMIGRATION SCENARIO | 32.1-12.3 | Dispersal | NSM | X | X | X |  | X |  |
| Hispaniolan *Poecilia* (Poeciliidae) – SEQUENTIAL IMMIGRATION SCENARIO | 32.1-12.3 | Dispersal | NSM |  | X |  |  |  |  |
| *Limia* (Poeciliidae) – SEQUENTIAL IMMIGRATION SCENARIO | 22.8-16.2 | Dispersal | NSM | X | X | X |  | X |  |
| *Rivulus berovidesi*-*R. cylindraceus* (Rivulidae) | ? | ? | NSM | X |  |  |  |  |  |
| *Rivulus roloffi* (Rivulidae) | ? | ? | NSM |  | X |  |  |  |  |
| *Cubanichthys* (Cubanichthyidae) | ? | Dispersal | NGM | X |  | X |  |  |  |
| **Neogene (23.0-2.6 Ma)** |  |  |  |  |  |  |  |  |  |
| *Gambusia melapleura*-*G. wrayi*-*G. hispaniolae* (Poeciliidae) | ~10.5 | Either | MT, WGM |  | X | X |  |  |  |
| *Gambusia rhizophorae*-*G. punctata*-(*G. beebei*-*G. pseudopunctata*-*G. xanthosoma*?<sup>1</sup>) (Poeciliidae) | ~8.7 | Dispersal | WGM | X |  |  |  | X |  |
| *Atractosteus tristoechus* (Lepisosteidae) | ? | Dispersal | NGM | X |  |  |  |  |  |
| *Gambusia manni*-*G. hubbsi* (Poeciliidae) | ~5.5 | Dispersal | MT |  |  |  |  |  | X |
| *Gambusia puncticulata*-*G. caymanensis*-*G. oligosticta* (Poeciliidae) | ~3.4 | Dispersal | MT | X |  | X |  | X | X |
| **Quaternary (2.6 Ma-present)** |  |  |  |  |  |  |  |  |  |
| *Cyprinodon bondi*-*C. nichollsi*-*C. sp. Lake Enriquillo*-(*C. laciniatus*?<sup>2</sup>) (Cyprinodontidae) | 2.2-1.7 | Dispersal | WGM, <u>NGM</u> |  | X |  |  |  | X? |
| *Cyprinodon variegatus* (Cyprinodontidae) | 0.8-0.4 | Dispersal | WGM, NSM, NGM | X | X | X |  | X | X |
| *Kryptolebias marmoratus* (Rivulidae) | 0.6-0.2 | Dispersal | NSM, EP | X |  |  |  | X | X |
| *Kryptolebias hermaphroditus* central clade (Rivulidae) | 0.5-0.1 | Dispersal | NSM, EP | X | X |  | X |  | X |
| *Fundulus grandis* (Fundulidae) | recent | Dispersal | NGM | X |  |  |  |  |  |

<sup>1</sup> Based on morphology, *G. xanthosoma* (Cayman Islands), *G. beebei*, and *G. pseudopunctata* (both from Hispaniola) are closely related to *G. rhizophorae* (Rivas, 1969)
<sup>2</sup> Phylogenetic placement is unknown for *C. laciniatus* (Bahamas), distinctive morphology (Richards & Martin, 2017) suggests it could represent an older immigration

As an alternative to long-distance dispersal, researchers have invoked a few landspan hypotheses to explain immigrations of freshwater fishes to the Greater Antilles. Rivas (1958) argued for a Yucatán-Cuba landspan to explain *Girardinus* (Poeciliidae) in Cuba because this genus is restricted to freshwater habitats. Ultimately, this hypothesis failed because geological evidence indicated Cuba was drifting away from Yucatán by 56 Ma (Pindell et al., 2005) and the Cuban lineage *Girardinus-Quintana* is not that old (Reznick et al., 2017; Table 1). Further, salinity tolerance is widespread throughout Poeciliidae, with many inland populations retaining substantial tolerance (Rosen & Bailey, 1963; Meffe & Snelson, 1989). Most Antillean poeciliids inhabit a range of habitats with varied salinities (Burgess & Franz, 1989; Haney & Walsh, 2003; García-Machado et al., 2020). Further, within Poeciliidae, offspring from the same parents may vary in salinity tolerance, which maintains potential for rapid adaptation from freshwater to a marine environment, or vice versa (Shikano & Fujio, 1997). This collective evidence demonstrates that a freshwater distribution may be insufficient, without additional evidence, to infer either salinity tolerance or biogeographic history within Poeciliidae.

Other landspan hypotheses (discussed below) are primarily supported as relevant for freshwater fishes by timing of estimated divergence in individual clades (e.g., Hrbek, Seckinger, & Meyer, 2007; Říčan et al., 2013; Weaver et al., 2016a). However, given that vicariance hypotheses predict the presence of replicate divergence events among unrelated taxa (Rosen, 1975; Platnick & Nelson, 1978), a synthesis of phylogenetic evidence among groups is needed to investigate collective patterns. Here, we assess hypotheses of immigration to the Greater Antilles with a literature review and synthesis of phylogenetic data for the entire freshwater-fish fauna. First, we summarize evidence for the three competing hypotheses of faunal assembly. Second, we conduct a regionalization of the fish fauna to delineate bioregions based on faunal similarity, providing context for biogeographical analysis. Third, we analyse possible patterns of immigration into the Greater Antilles bioregion, using the coefficient of dispersal direction and a minimum-spanning tree. Fourth, we conduct an ancestral-area reconstruction for each family including immigrants to the Greater Antilles. Fifth, for each immigrant lineage, we assess evidence bearing on whether a land connection or oceanic dispersal has most support. Where available, we summarize divergence-time estimates in relation to competing hypotheses. This includes a collective summary of all dated immigrations. Regarding terminology, we follow Crews & Esposito (2020) and eschew the term ‘colonization’ due to its imperial connotations. In addition to the term ‘dispersal’ (which has numerous and inconsistent usages; Wiley & Lieberman, 2011), we use the terms ‘immigration’, ‘establishment’, and ‘populate’.

## II. HYPOTHESES OF GREATER ANTILLEAN IMMIGRATIONS

### (1) Cretaceous vicariance hypothesis

The Cretaceous vicariance hypothesis postulates that landmasses in the Great Arc of the Caribbean (i.e., proto-Antilles) received immigrants from South America during tectonic collision, dated 100-90 Ma (Boschman et al., 2014; Stanek et al., 2019). However, evidence for intraplate collisions does not guarantee there were land connections (Perfit & Williams, 1989), and sea levels in the Cretaceous were higher than at present (Haq, 2014), decreasing potential for land connections. As an example, the Great Arc later collided with the Bahamas Platform (Eocene, Boschman et al., 2014), but land connections did not arise between the Greater Antilles and North America (Iturralde-Vinent, 2006), even though Cuba was transferred onto the North American block (Stanek, Maresch, & Pindell, 2009). It is also uncertain how much land (if any) in the Greater Antilles was continuously emergent in the years between the Cretaceous collision with South America and Eocene uplift in the Greater Antilles (Iturralde-Vinent, 2006; Roncal et al., 2020).

Early formulations of the Cretaceous vicariance hypothesis (Rosen, 1975; Burgess & Franz, 1989) preceded evidence that an asteroid collided with Earth on the Yucatán Peninsula, contributing to worldwide Cretaceous/Tertiary extinctions (Hildebrand et al., 1991). Well-established evidence for asteroid impact at Chicxulub ∼65.5 Ma now compromises the Cretaceous vicariance hypothesis because if any emergent land existed in the Greater Antilles when the collision occurred, proximate impacts (tsunami, wildfire) were likely catastrophic (Morgan et al., 2022; Santa Catharina et al., 2022). Substantial ejecta deposition extended >4,000 km from the impact site (Morgan et al., 2022), far enough to affect all Greater Antillean islands in their present positions. However, at 65.5 Ma, lands now forming Jamaica, Hispaniola, and Puerto Rico were west of their present positions, closer to Cuba and Yucatán (Boschman et al., 2014). Lands <500 km from the impact site, including areas in present-day Haiti (Hispaniola) were blanketed by dozens of meters of ejecta-rich deposits (Beloc formation, Hildebrand & Boynton, 1990; Schulte et al, 2010). Given that longer-range impacts ravaged distant terrestrial biotas (e.g., Lyson et al., 2019), faunas of the proto-Antilles were presumably devastated.

### (2) GAARlandia hypothesis

The GAARlandia (Greater Antilles-Aves Ridge) hypothesis proposes that 35-33 Ma, a peninsula, chain of islands, or combination extended from northern South America, along the Aves Ridge, to Puerto Rico, Hispaniola, east-central Cuba, and possibly eastern Jamaica (Iturralde-Vinent & MacPhee, 1999; Iturralde-Vinent, 2006). This hypothesis relies on evidence for Late Eocene uplift of the Aves Ridge, concurrent with sea-level fall (MacPhee & Iturralde-Vinent, 2005). The Great Arc did experience Late Eocene uplift, creating the GrANoLA landmass (Greater Antilles-Northern Lesser Antilles, Philippon et al., 2020). However, uplift of the Aves Ridge ended much earlier, ∼59 Ma (Late Palaeocene, Allen et al., 2019), and Aves Ridge has subsided since 52 Ma (Garrocq et al., 2021). Remaining physical evidence for GAARlandia is a two-step sea-level fall at ∼33.9 Ma (EOT-1, Eocene-Oligocene Transition 1) and ∼33.7 Ma (Oi1, Oligocene 1), totalling ∼75 m (dates and notations follow Miller et al., 2020a, acronyms indicate δ^18^O maxima, i.e., glacial periods). This may have been inadequate to expose the subsiding Aves Ridge, especially given that uplift is critical to the GAARlandia hypothesis (MacPhee & Iturralde-Vinent, 2005). There is evidence that submerged islands maintained coral reefs where coral growth kept pace with subsidence (Garrocq et al., 2021), in which case isolated reefs and possibly island tops were potentially exposed during sea-level fall.

Iturralde-Vinent & MacPhee (1999) devised the GAARlandia hypothesis with evidence from mammals, but subsequent analysis has diminished this support (Dávalos & Turvey, 2012; Woods et al., 2018, 2021). Primary supporting evidence now comes from fossils of Caribbean sloths (Delsuc et al., 2019) and caviomorph rodents (Marivaux et al., 2020), both of which also dispersed overseas. Sloths populated Central America prior to closure of the Isthmus of Panamá (Woodburne, 2010) and caviomorph rodents dispersed among Antillean islands (Woods et al., 2021). Relatives of anurans that potentially used GAARlandia (Alonso, Crawford, & Bermingham, 2012) also dispersed overseas (Heinicke, Duellman, & Hedges, 2007).

Several studies of terrestrial arthropods have indicated support for GAARlandia. Nevertheless, in a rare comparative study, Crews & Esposito (2020) found dispersal ability to be the best predictor of immigration to islands. Of 31 mainland-to-island dispersals, two (6.5%) were potentially associated with GAARlandia. One of these was later overturned (*Micrathena* spiders, Shapiro, Binford, & Agnarsson, 2022) and the remaining lineage (*Calisto* butterflies) only supported GAARlandia in one of two possible phylogenetic arrangements (Crews & Esposito, 2020). In other studies, *Deinopis* spiders potentially used GAARlandia, but at other times dispersed overseas (Chamberland et al., 2018). Three of five immigrations by Euophryinae spiders in the study of Zhang & Maddison (2013) possibly used GAARlandia, but one of these was subsequently revised with a younger date, incompatible with GAARlandia (Cala-Riquelme et al., 2022). For huntsmen (Sparassidae, Araneae), two immigrations potentially corresponded with GAARlandia, but two others occurred later, overseas (Tong et al., 2019).

Iturralde-Vinent & MacPhee (1999) proposed that evidence for vicariant speciation among Antillean islands supports the GAARlandia hypothesis, but vicariance among these islands could also occur for lineages established via overseas dispersal. Also, the GAARlandia hypothesis is vague regarding whether the proposed landspan was continuous or a chain of islands (Iturralde-Vinent & MacPhee, 1999; Iturralde-Vinent, 2006). This distinction is important because it is only possible to distinguish a landspan from long-distance dispersal if the land connection allowed range expansions by organisms otherwise incapable of overseas dispersal (Simpson, 1940; Hedges, 2006). If the GAARlandia landspan is treated as an archipelago rather than a landspan, then expectations are more similar (if not identical) to expectations from a hypothesis of overseas dispersal. Similarly, some authors have proposed that GAARlandia served as a dispersal route between South and Central America (Peña, Nylin, & Wahlberg, 2011; Říčan et al., 2013; Murillo-Ramos et al., 2021). Because GAARlandia could not have had a land connection to Central America (Iturralde-Vinent & MacPhee, 1999; Iturralde-Vinent, 2006), this scenario implies animals involved conducted overseas dispersal, and thus provides limited support for GAARlandia.

### (3) Null model for Antillean freshwater fishes

In the studies reviewed above, indicating supported for GAARlandia, there is an issue of parsimony because multiple mechanisms of immigration are inferred where one is sufficient. A simpler interpretation is that long-distance dispersal is the general mode of immigration to the Greater Antilles (Roncal et al., 2020). Another way to state this is that, without independent data for a landspan, and given evidence that the lineages in question can immigrate overseas, long-distance dispersal is an appropriate null hypothesis (Dávalos & Turvey, 2012). For instance, immigration timing was potentially congruent with GAARlandia for one lineage of amphisbaenids, but the preferred model for immigration in this group supports overseas dispersal, and the history of Amphisbaenia included two other overseas dispersals into the Caribbean (Graboski et al., 2022). For *Heraclides* swallowtails, although divergence estimates overlapped with the GAARlandia time frame, Lewis et al. (2015) documented two subsequent mainland-island immigrations and preferred a hypothesis of overseas dispersal. Further, studies document overseas immigration as the primary (or sole) mode for various Caribbean land mammals (Woods et al., 2018, 2021), lizards (Pinto-Sánchez et al., 2015; Tucker et al., 2017; Schools, Kasprowicz, & Hedges, 2022), ants (Price et al., 2022), weevils (Zhang et al., 2017), praying mantises (Svenson & Rodrigues, 2017), wasps (Ceccarelli et al., 2013; Rodriguez, Pitts, & von Dohlen, 2015), spiders (Agnarsson et al., 2016; Čandek et al., 2019, 2021), and land snails (Uit de Weerd, Robinson, & Rosenberg, 2016).

Given uncertainties associated with Cretaceous vicariance and a lack of geological evidence for GAARlandia, we believe it most parsimonious to treat dispersal as the null hypothesis for immigrations by freshwater fishes to the Greater Antilles. Neither landspan hypothesis includes geological evidence of freshwater connections between South America and the Greater Antilles. Absence of primary freshwater fishes from the Greater Antilles corroborates this evidence. In other words, reciprocal illumination supports an interpretation of long-distance dispersal for freshwater fishes immigrating to the Greater Antilles. Researchers invoking landspan hypotheses for freshwater fishes of the Greater Antilles rarely address the need for freshwater connections, but Weaver et al. (2016a) noted that if freshwater connections existed along GAARlandia, primary freshwater fishes should have reached the Greater Antilles. Notably, the Aves Ridge narrows to ∼30 km in width adjacent to South America (Garrocq et al., 2021). Even if this narrow isthmus emerged during Eocene-Oligocene sea-level fall, there might still have been an ocean gap between South America and wider portions of the Aves Ridge, farther north. Freshwater connections are often limited on a narrow peninsula or isthmus. For instance, Caribbean rivers of Costa Rica and Panamá are steep and short (Sosa Gonzalez et al., 2016), discharging onto a narrow continental shelf, suggesting little interconnection during sea-level falls (Bagley, Hickerson, Johnson, 2018). Accordingly, secondary freshwater fishes dominate the Central American fish fauna (Matamoros et al., 2014) because the Isthmus of Panamá restricted dispersal by primary freshwater fishes (Smith & Bermingham, 2005).

Additional examples are available. The Baja Peninsula has existed for >6 Ma (Umhoefer et al., 2018), twice as long as proposed for GAARlandia, but harbours no primary freshwater species (Miller, Minckley, & Norris, 2005; Rico et al., 2022). This is partly attributable to aridity, but also reflects limited riverine connectivity along the peninsula and across the narrow continental shelf (Dolby et al., 2018). The Florida and Yucatán peninsulas have broad continental shelves, but were subjected to Quaternary marine inundations. This, combined with low freshwater connectivity, produced depauperate freshwater-fish faunas, with few primary freshwater fishes (Gilbert, 1987; Miller et al., 2005; Elías et al., 2020). Thus, immigration along an isthmus or peninsula lacking direct freshwater connections appears to be restricted to the same species best suited for oceanic dispersal. This suggests that even if the Great Arc of the Caribbean or GAARlandia provided land bridges, without freshwater connections, they may not have facilitated immigration of freshwater fishes any more than an ocean gap would.

### (4) Considering ocean currents

Because of an historical emphasis on vicariance, syntheses of biogeography in Caribbean freshwater fishes often focus on changing land connections (Rosen, 1975; Burgess & Franz, 1989; Chakrabarty & Albert, 2011). However, if oceanic dispersal is an important mode of immigration, studies should also consider the historical geography of ocean currents. For instance, phylogeography of *Kryptolebias* in the Antilles, a presumed oceanic disperser, corresponds with the geography of contemporary currents (Tatarenkov et al., 2017). The dominant current system in the study region is the Caribbean-Loop current system (Fig. 3), flowing east to west across the Caribbean Sea, northwest through the Pedro and Yucatán channels to the Gulf of México, where it turns clockwise, southeast to the Straits of Florida, east through those straits, and then north along the Atlantic Coast of North America (Kirillova et al., 2019; Xu et al., 2022). This current system is an important route for intermittent rafting of flotsam and associated organisms (Thiel & Haye, 2006).

**Table 2.** Results of coefficient of dispersal direction (DD_2_) from mainland to the Greater Antilles bioregions. M = McNemar value. P = McNemar probability adjusted with the Holm correction (**** P < 0.0001).

| Bioregions | $DD_2$ | M | P |
| --- | --- | --- | --- |
| Northern Gulf of México (NGM) | 0.114 | 70 | **** |
| Western Gulf of México (WGM) | 0.103 | 14 | 0.0001 |
| Maya Terrane (MT) | 0.140 | 30 | **** |
| Chortís Block (CB) | 0.084 | 13 | 0.0003 |
| Eastern Panamá (EP) | 0.000 | 8 | 0.0042 |
| Northern South America (NSA) | 0.083 | 178 | **** |

Important Cenozoic events affecting the Caribbean-Loop current began with closure of the Georgia Channel in the Early Oligocene (Missimer, 2002; Missimer & Maliva, 2017). This directed outflow from the Gulf of México to the Straits of Florida (Guertin, Missimer, & McNeill, 2000) and, combined with narrowing of the Central American seaway (Late Oligocene, Montes et al., 2012; McGirr, Seton, & Williams, 2021), to establish the Caribbean-Loop Current system (Hübscher et al., 2023). This system serves as a dispersal avenue from northern South America to the Bahamas, with Central America, North America, and Cuba as points in between (Gaspar et al., 2022). Also in the Oligocene, uplift of the Andes Mountains created a continental, subandean river system that discharged into the Caribbean from north-western South America (Hoorn et al., 2010; Wesselingh & Hoorn, 2011). Sediments from this river extend across the Grenada Basin (Garrocq et al., 2021), suggesting this river discharged flotsam into the Caribbean Current.

Subsequent (Middle Miocene) narrowing of the Central American Seaway (Montes et al., 2015; McGirr et al., 2021) intensified the Caribbean-Loop Current (Hübscher et al., 2023), and connected it with the Mississippi-Tennessee delta in the northern Gulf of México (Xu et al., 2022). This suggests potential delivery of flotsam from these rivers into the Loop Current as it turned toward Cuba. However, for a time at the start of the Late Miocene (11.5-9.5 Ma), strong inflows from the Pacific Ocean into the Central American Seaway blocked the Caribbean Current from reaching the Gulf of México (Kirillova et al., 2019). Thereafter, progressive shoaling of the Central American Seaway (Late Miocene) eventually re-integrated and strengthened the Carribbean-Loop Current system (Kirillova et al., 2019; Prabhat et al., 2022).

## III. Synthetic Analyses

### (1) Clustering analysis and bioregions

Our study area included all coastal drainages from the Florida Peninsula to Venezuela, the Lesser Antilles, Greater Antilles, Bahamas (excluding Bermuda), and Cayman Islands. Our sampling unit was level-6 HydroBASINs (Linke et al., 2019). Data sources for fish distributions were Greenfield & Thomerson (1997), Miller et al. (2005), Smith & Bermingham (2005), Khin-Pineda, Cano, & Morales (2006), Maldonado-Ocampo, Vari, & Usma (2008), Rodríguez-Olarte, Taphorn, & Lobón-Cervía (2009), Matamoros, Schaefer, & Kreiser (2009), Matamoros, Kreiser, & Schaefer (2012), Matamoros et al. (2015, 2016), and DoNascimiento et al. (2017). Data were also extracted from Fishbase (Froese & Pauly, 2019). Nomenclature followed Eschmeyer, Fricke, & van der Laan (2022).

To delineate bioregions, we constructed a presence/absence matrix of 1,237 species (466 genera, 56 families), distributed among 96 level-6 HydroBASINS, and calculated a distance matrix of Jaccard pairwise dissimilarities (*beta.jtu* command, *beta.pair* function, R package *betapart*, Baselga et al., 2020). This method measures proportional replacement of species between assemblages, uninfluenced by species richness (Hattab et al., 2015). To produce a dendrogram of faunal similarity among hydrobasins, we used an unweighted paired-group method with arithmetic mean (UPGMA; *hclust* function, R package *stats*; R Core Team, 2021). To test how well the dendrogram maintained pairwise distances in the matrix, we calculated a cophenetic correlation (Farris, 1969; R package *Vegan*, Oksanen et al., 2020). This measure ranges from 0-1, with values >0.9 denoting excellent fit (Farris, 1969). The resulting dendrogram (Fig. 1) was an excellent fit to the data matrix (cophenetic correlation coefficient = 0.922).

**Fig. 1.**
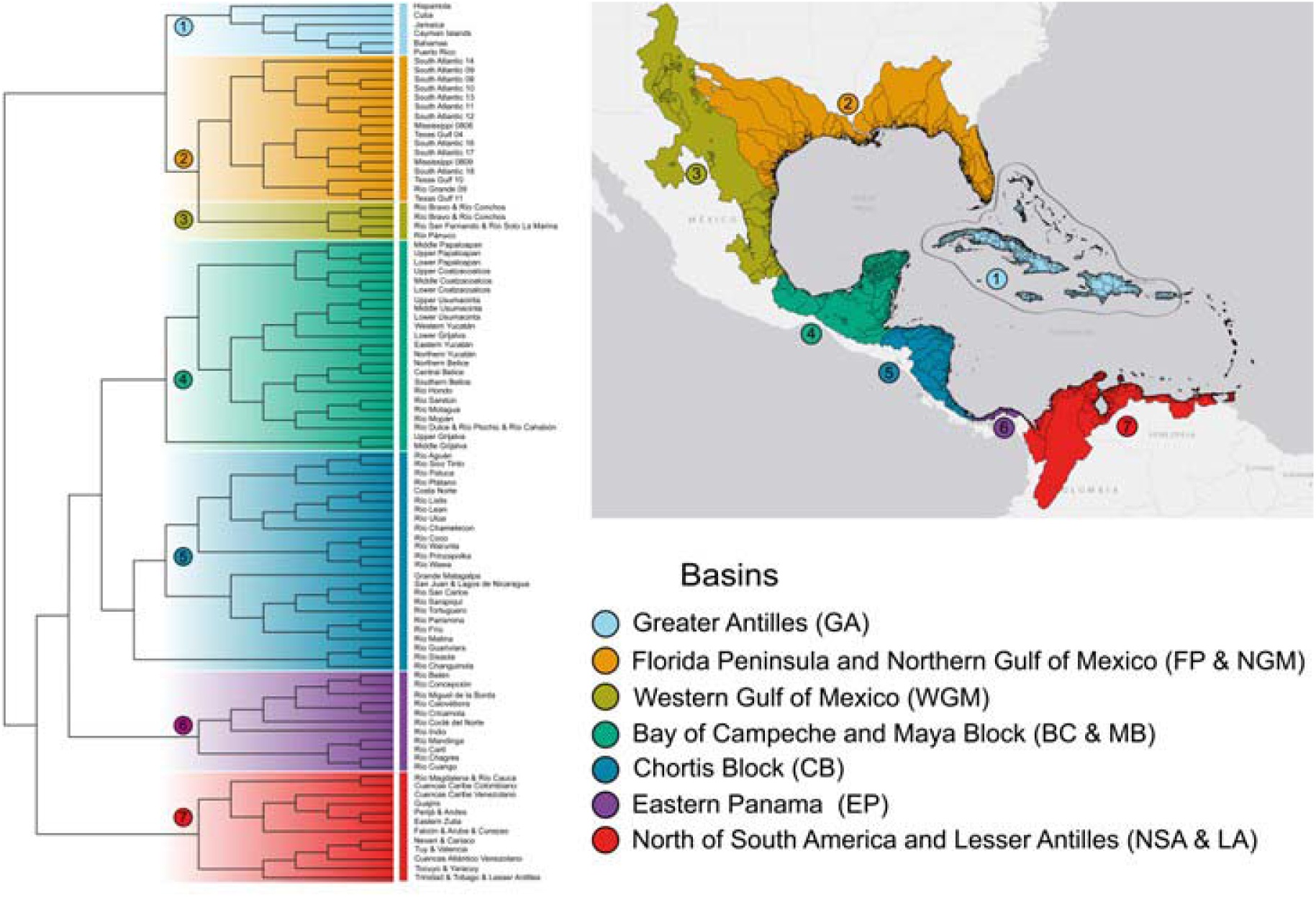
Colour coded beta diversity UPGMA (unweighted pair-group method using arithmetic averages) dendrogram of the Caribbean Sea and Gulf of México basins depicting 7 ichthyological bioregions (1) Greater Antilles (outlined), (2) Northern Gulf of México, (3) Western Gulf of México, (4) Maya Terrane, (5) Chortís Block, (6) Eastern Panamá, (7) northern South America (includes Lesser Antilles).

We applied the Kelley-Gardner-Sutcliffe penalty function (KGS; *kgs* function, R package *maptree*, White & Gramacy, 2015) to derive a minimum KGS value indicating the optimal number of clusters to maximize cohesiveness within branches (Kelley, Gardner, & Sutcliffe, 1996). The KGS function indicated 11 as the optimal number clusters on the UPGMA dendrogram. However, the tree had nine distinct branches, with five forks jointly positioned at the next level of branching (Fig. 1) forcing us to choose between 14 and 9 clusters. The 14-cluster scenario appeared to over-split the tree into small regions and even the 9-cluster scenario included two clusters with three or fewer HydroBASINs. One of these (upper-middle Grijalva) is a faunally distinct subdrainage (Elías et al., 2020), geographically nested within its sister branch, so we recognized this entire set of branches as the Maya Terrane bioregion (Fig. 1). In the other case, three HydroBASINs in western Panamá (Guariviara, Sixaola, Changuinola) formed a distinct branch sister to an adjacent region to the north. Rather than treat these western Panamanian HydroBASINs as a separate bioregion, we grouped them with their geographically contiguous sister branch (Fig. 1). From these considerations, we identified seven bioregions *(i)* Northern Gulf of México (NGM), *(ii)* Western Gulf of México (WGM), *(iii)* Maya Terrane (MT), *(iv)* Chortís Block (CB), *(v)* Eastern Panamá (EP), *(vi)* Northern South America (NSA, includes Lesser Antilles), and *(vii)* Greater Antilles (includes Bahamas and Cayman Islands, Fig. 1). To assess statistical validity, we investigated faunal similarity within and among branches using one-way analysis of similarity (ANOSIM, *anosim* function, *vegan* package in R, Oksanen et al., 2019). The resulting R test statistic ranges from 0 and 1 (*R* = 0, hydrobasins are similar among and within clusters; *R* = 1, hydrobasins within clusters are distinct from those in other clusters, Clarke & Warwick, 1994). ANOSIM results indicated the seven bioregion clusters were well differentiated (*R* = 0.78) and statistically different from each other (*P* < 0.001).

We documented 65 freshwater fish species (six families) in the Greater Antilles bioregion (Appendix S1). Because of the dominance of cyprinodontiforms in the Greater Antilles, we summarized the proportion of species from this family present in each bioregion as a measure of compositional dissimilarity. If immigration to the Greater Antilles was random for freshwater fishes, we would expect compositional similarity in proportions among bioregions. The pattern present (Fig. 2) indicates NSA and NGM bioregions have <10% cyprinodontiforms, with proportions increasing moving away from either continent, reaching 50% in the MT bioregion. Non-cyprinodontiform families from North America are diminished in the WGM and MT bioregions because deserts and mountains limit southward immigrations (Miller et al., 2005; Rico et al., 2022). Non-cyprinodontiform families from South America are diminished in the CB and MT bioregions due to barrier effects of the Isthmus of Panamá and former Central American Seaway (Matamoros et al., 2015; Bagley et al., 2018). Increased prevalence of cyprinodontiforms in the WGM, MT, and CB bioregions illustrates their superior dispersal ability. As biotic and environmental filtering can also block immigrants from becoming established after an island is reached (Schrader et al., 2021), cyprinodontiform adaptations may not only increase dispersal potential, but also facilitate establishment and persistence.

**Fig. 2.**
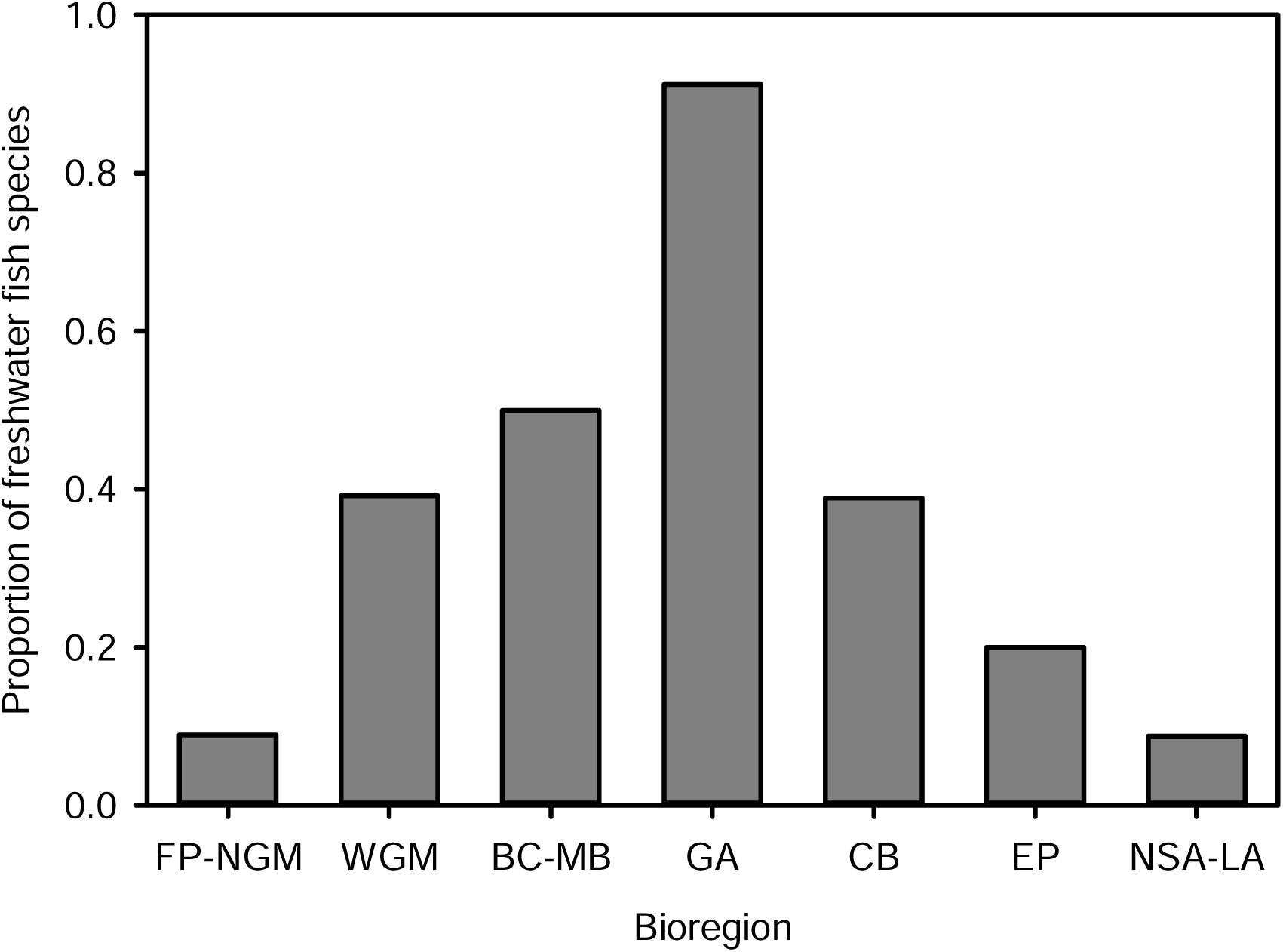
Proportion of fish species by bioregion that is within order Cyprinodontiformes. Species numbers by bioregion are: 281, Northern Gulf of México (NGM); 125, Western Gulf of México (WGM); 138, Maya Terrane (MT); 68, Greater Antilles (GA); 95 Chortís Block (CB); 65, Eastern Panamá (EP); 572, northern South America (NSA).

In comparison to a notable 50% cyprinodontiforms in the MT bioregion, the freshwater fauna of the Greater Antilles bioregion exceeds 90% cyprinodontiforms (Fig. 2). This illustrates that the Greater Antilles bioregion was even more difficult to reach and populate than the MT bioregion. Even Cichlidae, a prominent family of secondary freshwater fishes in Middle America (Matamoros et al., 2015), evidently had limited capacity to reach, persist, and diversify in the Greater Antilles. Hence, the freshwater-fish fauna of the Greater Antilles bioregion has exceptionally high compositional dissimilarity, in agreement with a hypothesis of long-distance dispersal.

### (2) Dispersal directions and minimum-spanning tree

To identify possible routes to the Greater Antilles, we used the coefficient of dispersal direction (DD_2_, Legendre & Legendre, 1984). The procedure makes four assumptions (Bachraty, Legendre, & Desbruye, 2009) *(i)* past dispersals leave traces in modern fish assemblages, *(ii)* dispersal is probable, *(iii)* dispersals occur only between neighbouring areas, and *(iv)* dispersals follow asymmetries, from areas with higher richness to areas with lower richness. We applied DD_2_,

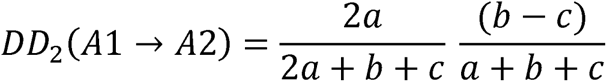

where *a* is the number of genera shared between two regions; *b* is the number of genera present in area 1 (A1) but not in area 2 (A2), and *c* is the number in A2 but not in A1 (we used the generic level to overcome the effect of local endemism in the Greater Antilles bioregion). The DD_2_ coefficient determines similarity between neighbouring faunas (first portion of formula, Sørensen index). Only faunas with taxa in common can be linked by dispersal. The coefficient also determines the direction of asymmetry (second portion of formula). In our analysis, if a genus immigrated from a mainland bioregion (A1) to the Greater Antilles (A2), the DD_2_ value would be positive, whereas if immigration occurred in the reverse direction (from A2 to A1), the DD_2_ value would be negative. For each DD_2_, we calculated the McNemar probability to verify asymmetry between bioregions (Bachraty et al., 2009; *bgdispersal* function, R package *vegan*, Oksanen et al., 2019). This analysis indicated richness asymmetry was present between the Greater Antilles and surrounding bioregions (Fig. 3a).

**Fig. 3.**
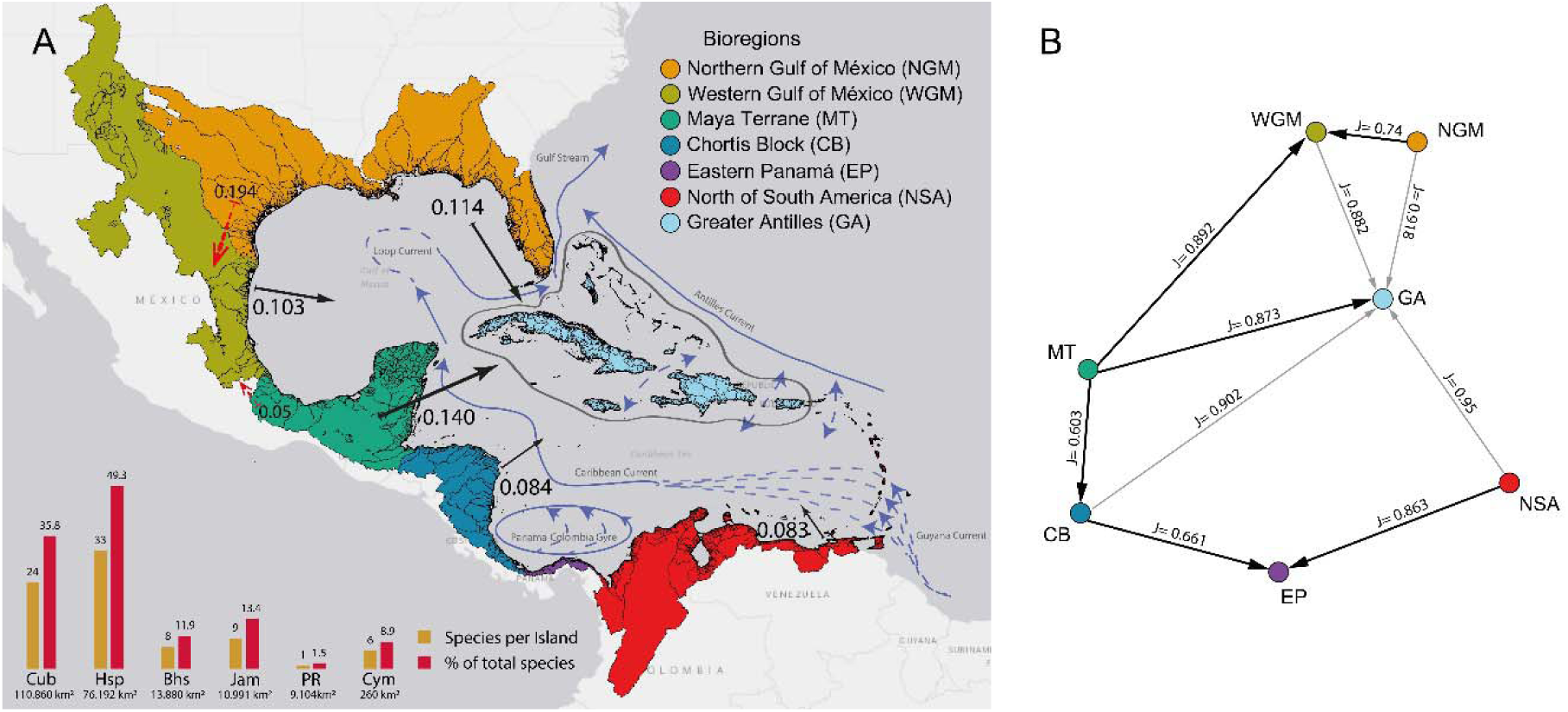
(A) Regionalization of 96 level-6 HydroBASINs of the Gulf of México and Caribbean Sea in a model with seven bioregions. Arrows indicate direction and magnitude (arrow size) of coefficients of dispersal direction (DD_2_) from mainland to the Greater Antilles (solid black lines) and among continental bioregions (dashed red lines). Species richness and percentage of total species richness in Greater Antilles with islands area are displayed at bottom left. In blue arrows, the marine currents that flows through the region. (B) Minimum spawning tree (MST, black arrowheads) among bioregions computed from Jaccard’s similarity coefficients using genera. Significant DD_2_ vectors (Table 2) that are not MST links are also shown (grey arrowheads). Jaccard’s values for generic similarity are displayed over the lines.

We used the Holm-Bonferroni sequential procedure to determine statistical significance of dispersal directions (Bachraty et al., 2009). Because the Greater Antilles bioregion has high faunal similarity to other bioregions (at the genus level), we used an alpha of 0.01 to identify bioregions with significant asymmetries. We also generated a minimum-spanning tree (MST) based on Jaccard similarity (Legendre & Legendre, 1998). Results were combined with information from the DD_2_ analysis to delineate faunal connections among bioregions. Based on strength of asymmetry (Table 2), the NGM, WGM, MT, CB, and NSA bioregions were likely source bioregions (Fig. 3b).

### (3) Ancestral-areas reconstruction

To reconstruct ancestral areas, we used published phylogenies when available. The Lepisosteidae phylogeny was based on a coalescent analysis of mitochondrial and nuclear genes (Wright, David, & Near, 2012). The Fundulidae phylogeny was generated from a 152-gene matrix constructed from RNA-sequence data for 16 species, combined with nuclear RAG1 and gylt gene sequences for 32 species (Rodgers et al., 2018). For Cichlidae, we used two topologies from Ilves, Torti, & López-Fernández (2018), which showed different positions for *Nandopsis* of the Greater Antilles, based on 415 exons for 139 species. Ilves et al. (2018) and Alda et al. (2021) recommended use of both topologies. The phylogeny for Poeciliidae was based on Reznick et al. (2017), which used nuclear and mitochondrial loci for 293 taxa. However, because Reznick et al. (2017) did not include Hispaniolan *Poecilia*, we also conducted an analysis for *Limia* and Hispaniolan *Poecilia* based on Weaver et al. (2016a).

Due to lack of published phylogenies for Cyprinodontidae and Rivulidae, we obtained cyt*b* sequences from GenBank for 24 species of Rivulidae and 21 species of Cyprinodontidae. These were aligned using the software Mega v.10.2 (Tamura et al., 2013). Ambiguities were coded as missing data. Bayesian inference analyses were performed using Mr. Bayes 3.2.7 on the alignments for each family (Ronquist & Huelsenbeck, 2003). Markov chain Monte Carlo was run for one million generations for two independent runs. We inspected the output for convergence and measured the average standard deviation of these runs until that value was <0.005, indicating the chains reached convergence.

We used ancestral-character-state reconstruction (Cunningham, Omland, & Oakley, 1998) to determine likely ancestral areas of taxa in each phylogenetic tree (Pirie et al., 2012). Each species was coded for its presence by bioregion and the procedure calculated possible ancestral areas for each node using parsimony in Mesquite 3.7, with minimum area change for seven areas (Pirie et al., 2012; Reznick et al., 2017). Ancestral-areas reconstructions indicated freshwater fishes of the Greater Antilles descend from 16 or 17 immigrations (discussed below). All adjacent bioregions potentially contributed immigrants to the Greater Antilles and each island group within the Greater Antilles bioregion harbours taxa from multiple ancestral areas, except for Puerto Rico, which supports only one species of freshwater fish (Table 1).

Evidence from the UPGMA tree (Fig. 1) and MST (Fig. 3b) indicated fish faunas are differentiated among three meta-regions: North America (NGM, WGM), Central America (MT, CB), and South America (EP, NSA). For an assessment of faunal filtering, we plotted the maximum number of immigrations from each meta-region to each island group of the Greater Antilles (Whittaker & Fernández-Palacios, 2007). Each meta-region contributed immigrants to each island group (except Puerto Rico). This pattern (Fig. 4) is counter to expectations of vicariance (Platnick & Nelson, 1978) and reveals a multi-way filtering effect, characteristic of island faunas assembled through long-distance dispersal (Whittaker & Fernández-Palacios, 2007).

**Fig. 4.**
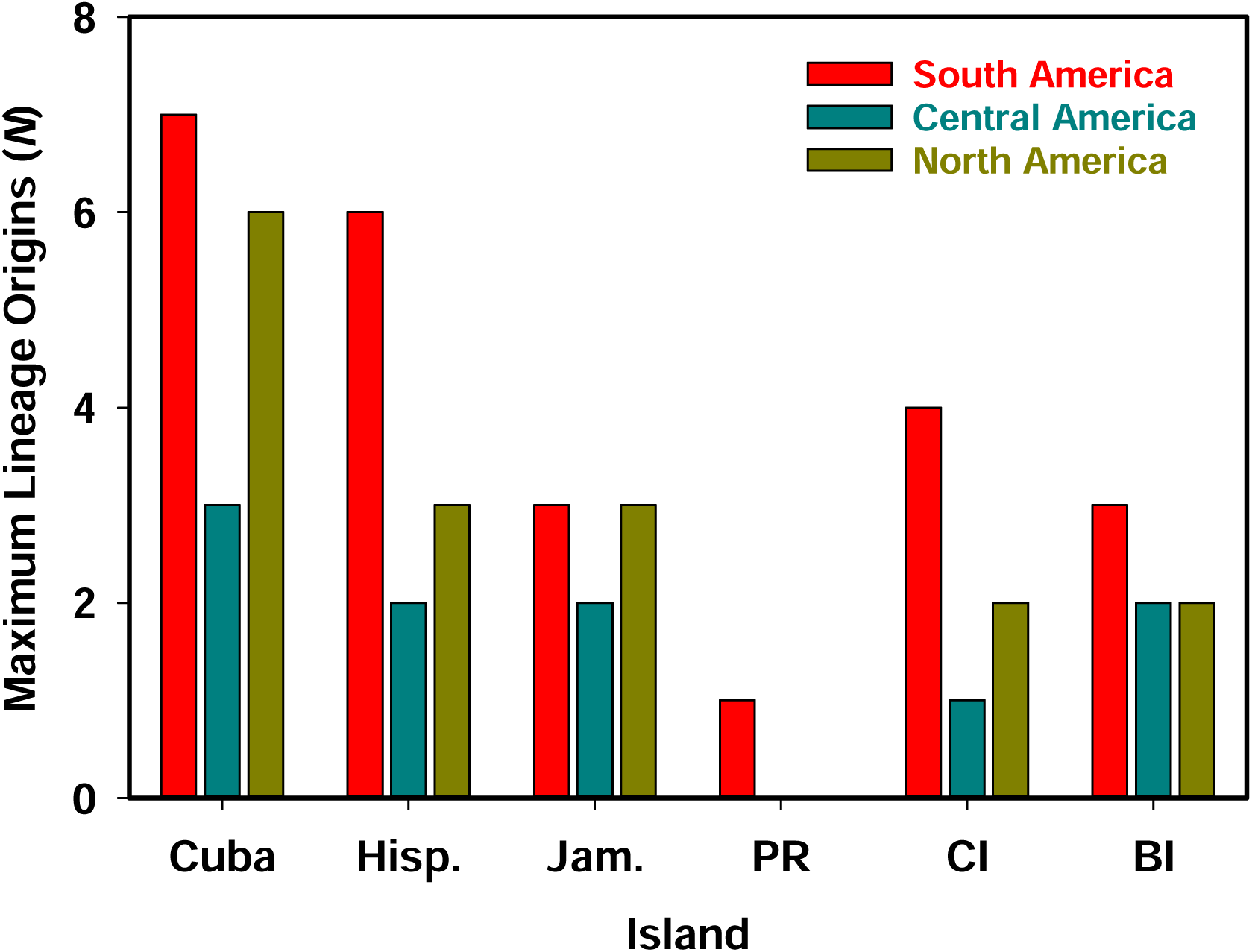
For each lineage of freshwater fish on each island group of the Greater Antilles bioregion, the maximum number of immigration events from each meta-region was summed. Meta-regions: South American (NSA, EP), Central American (CB, MT), North American (WGM, NGM). Taxa with multiple possible ancestral areas were counted for each continent that was a possible ancestral area (i.e., taxa could be counted multiple times).

### (4) Immigration narrative

We reviewed phylogenetic, geologic, and climatic literature, to interpret the succession of immigrations while assessing agreement with analytical findings. Because methods of interpretation vary, it is important to clarify our approach. We consider divergence estimates (i.e., nodes) to document timing of geographical separation between sister lineages (e.g., Weaver et al., 2016a). Some studies have used inter-node branch lengths to estimate separation timing (e.g., Reznick et al. 2017), but this implies divergence preceded geographical separation, in contrast to the assumption of vicariance biogeography, that geographical separation causes divergence (Rosen, 1975; Platnick & Nelson, 1978). To support a vicariance scenario, confidence limits of a divergence estimate should overlap with the timing of barrier formation (e.g., Alonso et al., 2012; Delsuc et al., 2019). Some studies appear to dismiss confidence limits in favour of a landspan hypothesis, despite incongruent timing (Říčan et al., 2013; Deler-Hernández et al., 2018; Cala-Riquelme et al., 2022), but we used a stricter approach. For any reference cited, we use reported divergence estimates, not inter-node lengths. If a confidence estimate did not overlap in time with timing of land separation, we concluded that a relation of phylogenetic divergence with landspan disruption was improbable. Immigrations are narrated below in chronological order. For undated immigrations, the order is tentative based on available information.

#### (a) Paleogene (66.0-23.0 Ma)

i. Girardinus-Quintana. The ancestral area for *Girardinus-Quintana* (Poeciliidae) is ambiguous and could be the WGM, MT, EP, or NSA bioregions (Fig. 5). Although *Girardinus-Quintana* has the oldest divergence estimate available for freshwater fishes of the Greater Antilles (Table 1), and the 95% credibility intervals are broad (50.7-22.3 Ma) and provisional (Reznick et al., 2017), the divergence estimate greatly post-dates the 100-90 Ma Cretaceous vicariance scenario, in disagreement with Doadrio et al. (2009). The credibility intervals do overlap with the Oi1 glaciation, which technically supports the GAARlandia hypothesis, in agreement with Hrbek et al. (2007) and Reznick et al. (2017). However, the intervals also overlap with other sea-level rises and falls (Fig. 6). The point estimate most closely aligns with the Middle Eocene Climatic Optimum, a time of sea-level rise. Without further information, this immigration remains mysterious.
ii. Nandopsis. *Nandopsis* (Cichlidae) represents one of only two immigrations to the Greater Antilles by non-cyprinodontiforms. A recent divergence estimate (Říčan et al., 2013) is too young for Cretaceous vicariance, in disagreement with Chakrabarty (2006; Chakrabarty & Albert, 2011). It is also at least 4.0 Ma younger than the Oi1 glaciation (GAARlandia), in disagreement with Říčan et al. (2013). At the latest, post-glacial sea-level rise at 32.0 Ma (Miller et al., 2020a) would have disrupted GAARlandia (if it existed), and this occurred at least 2.5 Ma before the estimated divergence of *Nandopsis* (Fig. 6). High salinity tolerances in Central American cichlids and presence in coastal and marine waters (Oldfield, 2007), along with discovery of a Late Miocene cichlid fossil in marine sediments of Costa Rica (Lucas et al., 2017), suggest potential for oceanic dispersal, in agreement with classifications of cichlids as secondary freshwater fishes (Myers, 1938; Bianco & Nordlie, 2008) and freshwater-estuarine opportunists (Potter et al., 2013; Whitfield, 2015). Further, a Cichlidae-Pholidichthys sister relationship (Eytan et al., 2015) suggests the MRCA of Cichlidae was marine (Matschiner et al., 2020). Divergence estimates and ancestral-area reconstructions suggest the MRCA of New World cichlids reached South America from Africa, across the open ocean (Lavoué, 2020). Thus, New World cichlids likely descend from an ocean-crossing ancestor. The phylogenetic position of *Nandopsis* is uncertain (Ilves et al., 2018) and trees used in this analysis (Fig. 7) indicate different placements than in Říčan et al. (2013). Thus, the applicability of the available divergence estimate is unclear. Following the Astral-II topology (Ilves et al., 2018), *Nandopsis* is the earliest diverging lineage from a clade that populated the Chortís Block, suggesting the MRCA was distributed among the NSA, CB, and Greater Antilles bioregions (Fig. 7). If the MRCA inhabited the subandean river, rafts of vegetation discharged into the Central American Seaway may have been carried, perhaps within a plume of brackish water (Tagliacollo et al., 2017), into the Central American Seaway. In the Oligocene, the seaway current flowed westward (Fraass et al., 2019). This would have transported the subandean-river plume toward the Pacific Ocean, but Late Eocene narrowing of the seaway as the Panamá Arc collided with South America (Montes et al., 2012; McGirr et al., 2021) could have increased potential for landfall en route. Once established, the MRCA of Chortís Block cichlids potentially dispersed along the Nicaragua Rise, which hosted continuous carbonate banks and barrier reefs extending to Jamaica (Roth, Droxler, & Kameo, 2000; Mutti, Droxler, & Cunningham., 2005). Previous studies suggest cichlids used this route in one direction or another (Chakrabarty & Albert, 2011; Říčan et al., 2013). In the Oligocene, most of Jamaica was submerged (Gold et al., 2018), but Hispaniola and Cuba (where *Nandopsis* occurs) were adjacent to the rise and partly emergent (Iturralde-Vinent, 2006), suggesting availability to cichlids. The carbonate banks on the Nicaragua Rise began to founder (i.e., sink, collapse) in the Late Oligocene, allowing the Caribbean Current to bisect the Nicaragua Rise ∼27 Ma (Mutti et al., 2005), potentially isolating Antillean populations from those on the Chortís Block, congruent with the available divergence estimate for *Nandopsis* (Fig. 6). In comparison, following the STAR topology (Fig. 7), *Nandopsis* is nested within South American cichlids, suggesting direct immigration to Hispaniola or Cuba from South America. The present *Nandopsis* divergence estimate precedes a period of current reversal in the Central American Seaway that temporarily allowed Pacific inflow into the Caribbean, ∼23 Ma (Fraass et al., 2019). This could have facilitated northward transport of rafts from the subandean river and aligns in time with the Mi1 sea-level fall (Miller et al., 2020a), which could have shortened distances from South America to Hispaniola and Cuba. If future phylogenies support the STAR topology, this scenario may deserve further consideration, but will also require a younger divergence estimate for strong support.
iii. Limia *+ Hispaniolan* Poecilia. *Limia* (Poeciliidae), with all but one species in the Greater Antilles, is sister to three Hispaniolan species of *Poecilia (P. dominicensis, P. hispaniolana, P. elegans*, Palacios et al., 2016; Weaver et al., 2016a). Our ancestral-area reconstructions indicate South America as the ancestral area (Fig. 5), implying direct immigration. This agrees with persistence of saltwater tolerance among *Limia* (Burgess & Franz, 1989; Haney & Walsh, 2003; Weaver et al., 2016b). Although Weaver et al. (2016a) found congruent timing of *Limia* divergence with GAARlandia, other studies produced younger estimates (Hrbek et al., 2007; Reznick et al., 2017). The estimate used here is provisional (Reznick et al., 2017), but suggests Oligo-Miocene immigration. Environmental factors that could have enhanced immigration include (*i*) Oligocene discharge from the subandean river, (*ii*) collision of the Panamá Arc with South America, invigorating the Caribbean Current, and (iii) Mi1 sea-level fall, shortening travel distance (Fig. 6). In addition, viviparity in *Limia* (Helmstetter et al., 2016) could increase rafting potential (Thiel & Gutow, 2005). It is uncertain whether *L. heterandria* of Venezuela arose before or after establishment of *Limia* in the Antilles. Either hypothesis requires two immigrations. If the MRCA for the entire *Limia-*Hispaniolan *Poecilia* clade immigrated to the Antilles, ancestral *L. heterandria* descends from an immigrant back to South America. Alternatively, if Hispaniolan *Poecilia* immigrated first, *Limia* in South America possibly diverged from Hispaniolan *Poecilia* in situ and later immigrated to the Greater Antilles, leaving *L. heterandria* behind. An ancestral-areas analysis based on a phylogeny that includes all taxa in this group may help resolve this uncertainty.
iv. Rivulus. Murphy & Collier (1996) inferred, without formal divergence estimates, that Antillean *Rivulus* (Rivulidae) descend from immigrations occurring 80-70 Ma, leading them to favour Cretaceous vicariance. This is questionable because, not only was the dating method informal, but Late Cretaceous immigration implies that Antillean *Rivulus* survived in proximity to the bolide impact (described above). There is also phylogenetic uncertainty as some reconstructions resolve *R. berovidesi*-*R. cylindraceus* (Cuba) and *R. roloffi* (Hispaniola) as sister taxa, but others resolve them as independent branches (Murphy, Thomerson, & Collier, 1999). Our analysis agrees with the latter scenario (Fig. 9), suggesting separate immigrations to each island from NSA. *Rivulus hartii* conduct oceanic dispersal (Walter et al., 2011) and exhibit egg diapause under harsh environmental conditions (Furness, 2016). If ancestors of Antillean *Rivulus* shared similar adaptations, this could have increased chances of oceanic immigration. Based on the phylogenetic positions of Antillean *Rivulus* (Fig. 9), we speculate that both immigrations occurred in the Paleogene (Table 1).
v. Cubanichthys*. Cubanichthys cubensis* (Cuba) and *C. pengelleyi* (Jamaica) are highly divergent (Lara et al., 2010), and *Cubanichthys* (Cubanichthyidae) may be paraphyletic (Piller et al., 2022). Further, the phylogenetic position of *Cubanichthys* varies among studies (Helmstetter et al., 2016), and the poorly known *Yssolebias martae*—a single specimen from the Río Magdalena delta, Colombia—may be related (Huber, 2015). These uncertainties preclude detailed biogeographical assessment. The origins of *Cubanichthys* are undated, but as the basal split within Cyprinodontidae was 23.8-18.4 Ma (Echelle et al., 2005), *C. cubensis* and *C. pengelleyi* presumably predate this. However, Jamaica was not emergent until 25 Ma or later (James-Williamson, Mitchell, & Ramsook, 2014), suggesting *C. pengelleyi* is younger than 25 Ma but older than 18.4 Ma, leading us to speculate a Late Oligocene-Early Miocene origin (Table 1). Ancestral area reconstruction indicates *C. cubensis* immigrated from the NGM bioregion (Fig. 10), but this result is tentative given uncertain phylogenetic placement.

#### (b) Neogene (23.0-2.6 Ma)

i. Gambusia. Reznick et al. (2017) identified four instances of *Gambusia* (Poeciliidae) reaching Caribbean islands. Viviparity likely provided an advantage for raft immigrations. The earliest immigration established *G. melapleura-G. wrayi-G. hispaniolae* of Jamaica and Hispaniola. Ancestral areas for this lineage could be the WGM or MT bioregions (Fig. 5). From 11.5-9.5 Ma, a window of time congruent with divergence of this lineage (Fig. 6), inflow from the Pacific Ocean blocked the Caribbean Current, and offshore currents from the MT bioregion were directed toward the Greater Antilles (Kirillova et al., 2019). The MRCA potentially reached Jamaica in stepwise fashion along the northern Nicaragua Rise, where remnant carbonate banks and barrier reefs persisted (Roth et al., 2000; Mutti et al., 2005). Flow of the Caribbean-Loop Current resumed through the Pedro and Yucatán channels ∼9.0 Ma (Kirillova et al., 2019) and ∼8.7 Ma, the *G. punctata* species group is estimated to have populated Cuba from the WGM bioregion (Figs. 5, 6). At this time, the ancestral ríos Grande and San Fernando extended to the Sierra Madre Oriental (Snedden & Galloway, 2019), which hosted tropical deciduous forests (Pound et al., 2011). These rivers potentially discharged rafts of vegetation into an offshore current, which joins the Loop Current where it turns toward Cuba (Sanvicente-Añorve et al., 2014; Xu et al., 2022) (Fig. 3.). Recent immigration of *G. rhizophorae* from Cuba to Florida (García-Machado et al., 2020) supports a history of oceanic dispersal. Thereafter, the MRCA of *G. manni-G. hubbsi* populated the Bahamas ∼5.5 Ma (Reznick et al., 2017). Our analysis identifies the MT bioregion as the ancestral area (Fig. 5) theoretically via the Loop Current. Later still, the *G. punticulata-G. caymanensis-G. oligosticta* lineage populated the Greater Antilles ∼3.4 Ma (Reznick et al., 2017), likely also from the MT bioregion (Fig. 5). This lineage inhabits Cuba, Jamaica, and the Cayman Islands, suggesting immigration through the Caribbean Sea as proposed for *G. melapleura-G. wrayi-G. hispaniolae* (above). However, divergence timing corresponds with the Pliocene Climatic Optimum (Fig. 6), a period of high sea level (Miller et al., 2020a), and post-dates establishment of the Caribbean-Loop Current through the Pedro Channel (Kirillova et al., 2019), suggesting instead that immigration was via the Loop Current.
ii. Atractosteus. *Atractosteus spatula* (sister to Cuban *A. tristoechus*, Lepisosteidae) is salt-tolerant (Grande, 2010; Echelle & Grande, 2014), sometimes found on the ocean side of barrier islands (Suttkus, 1963) or in the open ocean (Gunter, 1942). Although *Atractosteus* attaches eggs to vegetation, larvae and small juveniles have low salt tolerance (Echelle & Grande, 2014), suggesting larger juveniles or adults immigrated to Cuba. There is no divergence estimate for *A. tristoechus*. Wiley (1976) hypothesized that a shared biogeographical pattern with *Gambusia* suggested shared timing. Our analysis identified the NGM bioregion as the ancestral area for *A. tristoechus* (Fig. 10), which contrasts with *Gambusia* that immigrated from farther west. Fossil evidence of *Atractosteus* on the Florida Peninsula in the Pleistocene (Hay, 1919; Grande, 2010) opens the possibility that the genus dispersed across the Straits of Florida and persisted in Cuba while disappearing from southern Florida. If *Atractosteus* inhabited the Florida Peninsula in the Late Miocene-Pliocene, the Great Floridian River, which discharged into the Straits of Florida (Missimer & Maliva, 2017), might have facilitated immigration, as hypothesized for *Trachemys* turtles (Hoagstrom et al., 2022).

#### (c) Quaternary (2.6 Ma to present)

i. Cyprinodon. Hispaniolan *Cyprinodon bondi-C. nichollsi-C*. sp. Lake Enriquillo separated from North American relatives in the Early Pleistocene (Haney et al., 2007). Echelle et al. (2006) hypothesized immigration occurred during sea-level falls, several of which occurred in this period (Miller et al., 2020a). Hispaniolan pupfish belong to a maritime clade that ranges through the NGM, WGM, NSA, and Greater Antilles bioregions (Echelle et al., 2005, 2006). Our reconstruction identifies the WGM as the ancestral area (Fig. 10), but phylogeography indicates Atlantic Coast populations are the closest to Hispaniolan endemics (Echelle et al., 2006). This discrepancy reveals a limitation of ancestral-areas analysis. The maritime clade includes several disjunct peripheral isolates (Lozano-Vilano & Contreras-Balderas, 1999; Echelle et al., 2006; Haney, Turner, & Rand, 2009; Brix & Grosell, 2013), *C. tularosa* (WGM Chihuahuan Desert), *C. bobmilleri* (WGM Coastal Plain), *C. dearborni* (NSA), *C. bondi-C. nichollsi-C*. sp. Lake Enriquillo (Hispaniola), and *C. variegatus hubbsi* (NGM Florida Peninsula). Peripheral isolates diverge separately from the widespread stem lineage, not from each other, confounding ancestral-areas analysis. *Cyprinodon variegatus* is a widespread, euryhaline species, inhabiting the NGM, WGM, NSA, and Greater Antilles bioregions (Fig. 10). It is widespread among Caribbean Islands (Smith, Rodriguez, & Lydeard, 1990) and may include *C. higuey* of Hispaniola (Echelle et al., 2006; Haney et al., 2007). Initial immigration to the Greater Antilles at 0.76-0.40 Ma was likely associated with sea-level falls (Echelle et al., 2006; Haney et al., 2007). Phylogeographic evidence indicates there has been widespread gene flow throughout the Greater Antilles and Florida Peninsula (Echelle et al., 2006; Richards et al., 2021). Dramatic sea-level falls of the Middle-Late Pleistocene may have facilitated immigrations (Fig. 6).
ii. Kryptolebias. Two species of *Kryptolebias* (Rivulidae) inhabit the Greater Antilles (Berbel-Filho et al., 2022) (Fig. 9). Cuban and Bahaman *K. marmoratus* belong to a lineage also present on the Florida Peninsula and Florida Keys. Genetic evidence indicates *K. marmoratus* conducts oceanic dispersal (Tatarenkov et al., 2012, 2015). Potential for adhesive embryos to survive ocean salinity attached to flotsam and ability of a single selfing individual to found a new population likely enhance dispersal ability (Tatarenkov et al., 2017). Ability of juveniles and adults to survive out of water may also be advantageous (Turko & Wright, 2015). In any case, the Florida-Cuba-Bahamas lineage of *K. marmoratus* diverged from MT-CB populations 0.6-0.2 Ma (Tatarenkov et al., 2017), a time of intense, periodic glaciation and attendant sea-level fluctuation (Fig. 6). Also, the Loop Current is truncated during glacial periods, flowing more directly from the Yucatán Channel into the Straits of Florida (Arellano-Torres, Amezcua-Montiel, & Casas-Ortiz, 2023), which could complement sea-level falls in reducing travel distance. *Kryptolebias hermaphroditus* inhabits the Greater Antilles, NSA, EP, and CB bioregions (Fig. 9). Island immigration is dated 0.5-0.1 Ma (Tatarenkov et al., 2017). This species has a unique distribution pattern including the Lesser Antilles and Puerto Rico (Berbel-Filho et al., 2022), suggesting dispersal via the Antilles Current, as observed in juvenile leatherback turtles *Dermochelys coriacea* (Gaspar, Candela, & Shillinger, 2022). However, a population of *K. hermaphroditus* in EP-CB bioregions indicates the species also either dispersed via the Caribbean Current to the Panamá-Columbia Gyre or along the northern coast of South America.
iii. Fundulus. *Fundulus grandis* (Fundulidae) inhabits the WGM, NGM, and Greater Antilles bioregions (Fig. 9). Southern Florida and Cuban populations are described as *F. grandis saguanus* (Rivas, 1948; Relyea, 1983), but molecular evidence refutes this hypothesis for Floridian populations (Duggins, Relyea, & Karlin, 1989), casting doubt on the status of Cuban populations. Glaciations pushed *F. grandis* south into disjunct climatic refugia, one in the WGM bioregion and another on the Florida Peninsula (Williams, Brown, & Crawford, 2008). Ancestral-area reconstruction indicates either lineage could have immigrated to Cuba (Fig. 9). Nevertheless, proximity and morphology imply *F. grandis* reached Cuba from Florida (Relyea, 1983; Rivas, 1986). Rivas (1948) suggested Cay Sal Bank could have been a stepping stone. Consistent with this hypothesis, the 6000 km^2^ bank was 99% exposed 11 ka (Purkis et al., 2014). Like *C. variegatus*, *F. grandis* speciates via peripheral isolation (García-Ramírez, Contreras-Balderas, & Lozano-Vilano, 2006, García-Ramírez, Lozano-Vilano, & de la Maza Benignos, 2021). Hence, phylogeographic study is needed to determine taxonomy and ancestral area of Cuban *F. grandis*.

#### (d) Summary discussion

Cyprinodontiformes dominates the Greater Antilles freshwater fish fauna in immigration events (15 of 17 possible) and species richness (Fig. 2). Evidence indicates Cyprinodontiformes immigrated, sequentially (albeit rarely), and from various bioregions. This fits the dispersal paradigm, that organisms better adapted for challenges of oceanic dispersal, population establishment, and population persistence are most likely to accomplish it. We document a maximum of 15 cyprinodontiform immigrations within the last 51 my, one immigration every 3.4 my on average (although, widespread species with evidence of gene flow across the Greater Antilles and among neighbouring bioregions may have made more frequent immigrations). During this 51 my period, cichlids accomplished one immigration, possibly aided by the Nicaragua Rise before it foundered. Gars (Lepisosteidae) also appear to have accomplished just one immigration, possibly across the relatively narrow Straits of Florida. If any other group of fishes ever accomplished an immigration, descendants were evidently extirpated before documentation. This pattern is diametrically opposed that expected in vicariant speciation (Platnick & Nelson, 1978).

**Fig. 5.**
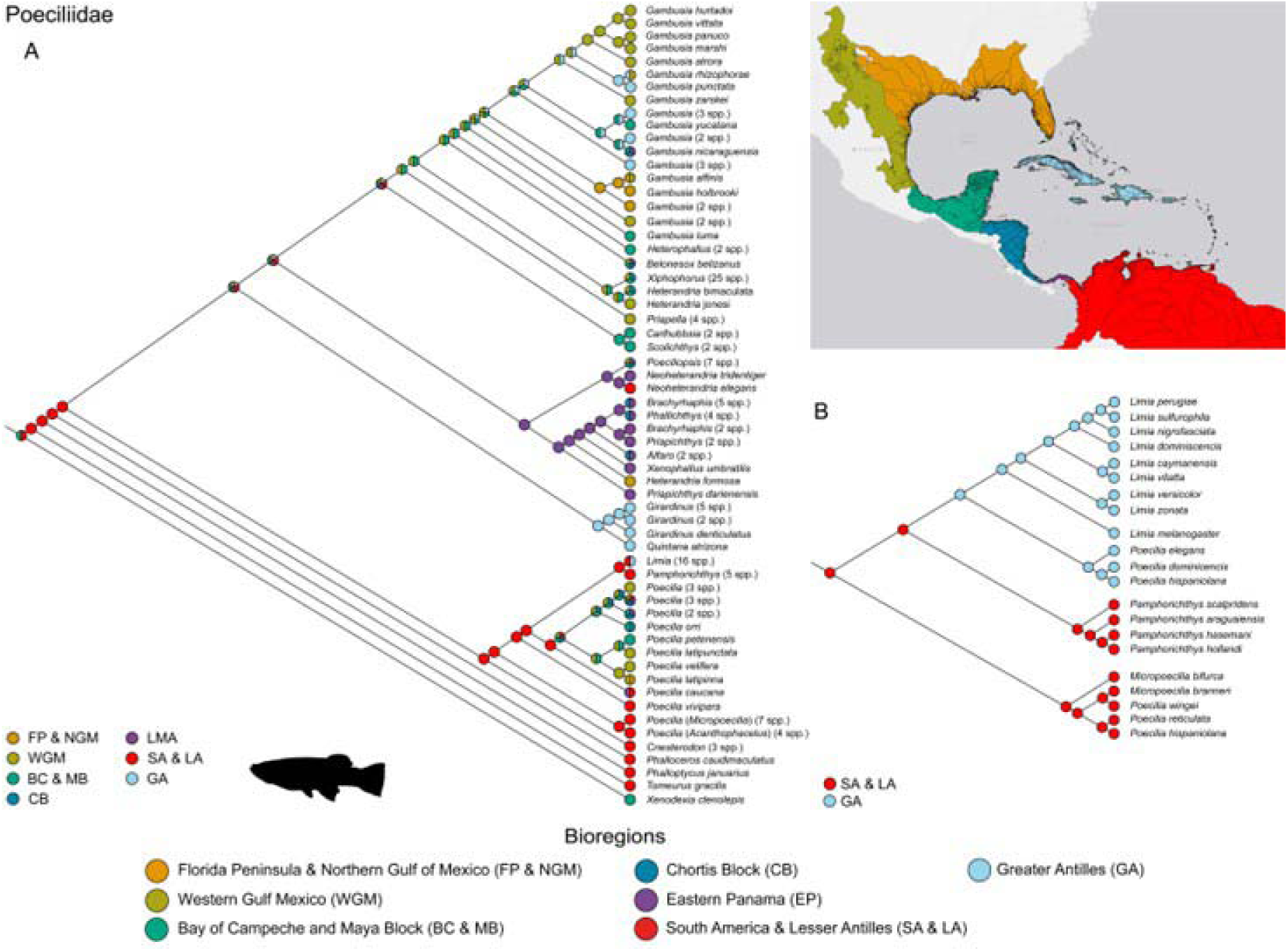
Final ancestral areas analysis redrafted from Reznick et al. (2017) for Poeciliidae (A). This tree is missing Hispaniolan Poecilia, so an analysis redrafted from Weaver et al. (2016a) is provided for Antillean *Limia* and *Poecilia* (B).

**Fig. 6.**
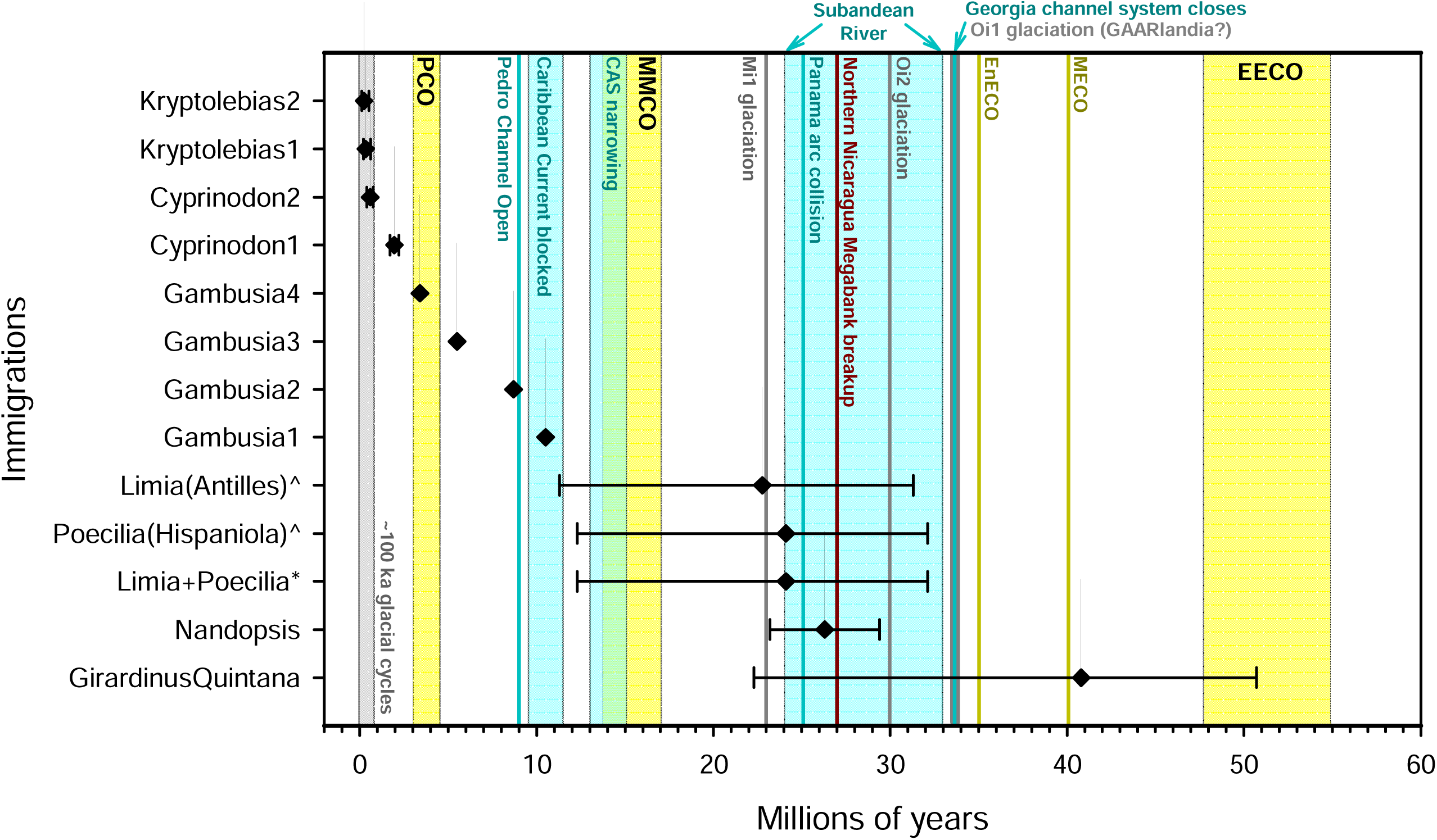
For immigrations that have estimated timing (Table 1), estimates are plotted on a timeline showing major oceanographic events (Miller et al. 2020a, 2020b). For *Gambusia*, only point estimates are available (Reznick et al., 2017). Other estimates include error bars indicating measures of uncertainty provided in references (Haney et al., 2007; Reznick et al., 2017; Tatarenkov et al., 2017; Pérez-Miranda et al., 2020). Two possible scenarios for *Limia* & Hispaniolan *Poecilia—*as a combined (asterisk) or as separate (caret) immigrations—are shown (see text). Warm periods with elevated sea levels are show in yellow (EECO – Early Eocene climatic optimum [55.0-47.9 Ma]; MECO – Middle Eocene climatic optimum [40.1 Ma]; EnECO – End Eocene climatic optimum [35.0 Ma], MMCO – Middle Miocene climatic optimum [17.0-13.8 Ma]; PCO – Pliocene climatic optimum [4.5-3.0 Ma]). Glacial periods with exceptionally low sea levels are shown in grey (EOT1-Oi1, 33.9-33.7 Ma; Oi2, 30.0 Ma; Mi1, 23.0 Ma; Middle-Late Pleistocene, 100 ka glacial cycles). Note, EOT1-Oi1 glaciations correspond in time with the GAARlandia hypothesis. Reorganization of ocean currents is shown in blue-green: closure of the Georgia channel system (33.7 Ma, Missimer, 2002); discharge of subandean river into the Granada Basin (33.0-24.0 Ma, Wesselingh & Hoorn, 2011); collision of Panamá arc with South America (25.0 Ma, Montes et al., 2012); Central American Seaway (CAS) narrowing (15.0-13.0 Ma, Montes et al., 2015); break in the Caribbean Current due to Pacific Water inflow (11.5-9.5 Ma, Kirillova et al., 2019), and establishment of the Caribbean Current through the Pedro Channel (9.0 Ma, Kirillova et al., 2019). Foundering of the Nicaragua Rise ∼27 Ma (Mutti et al., 2005) is shown in burgundy.

**Fig. 7.**
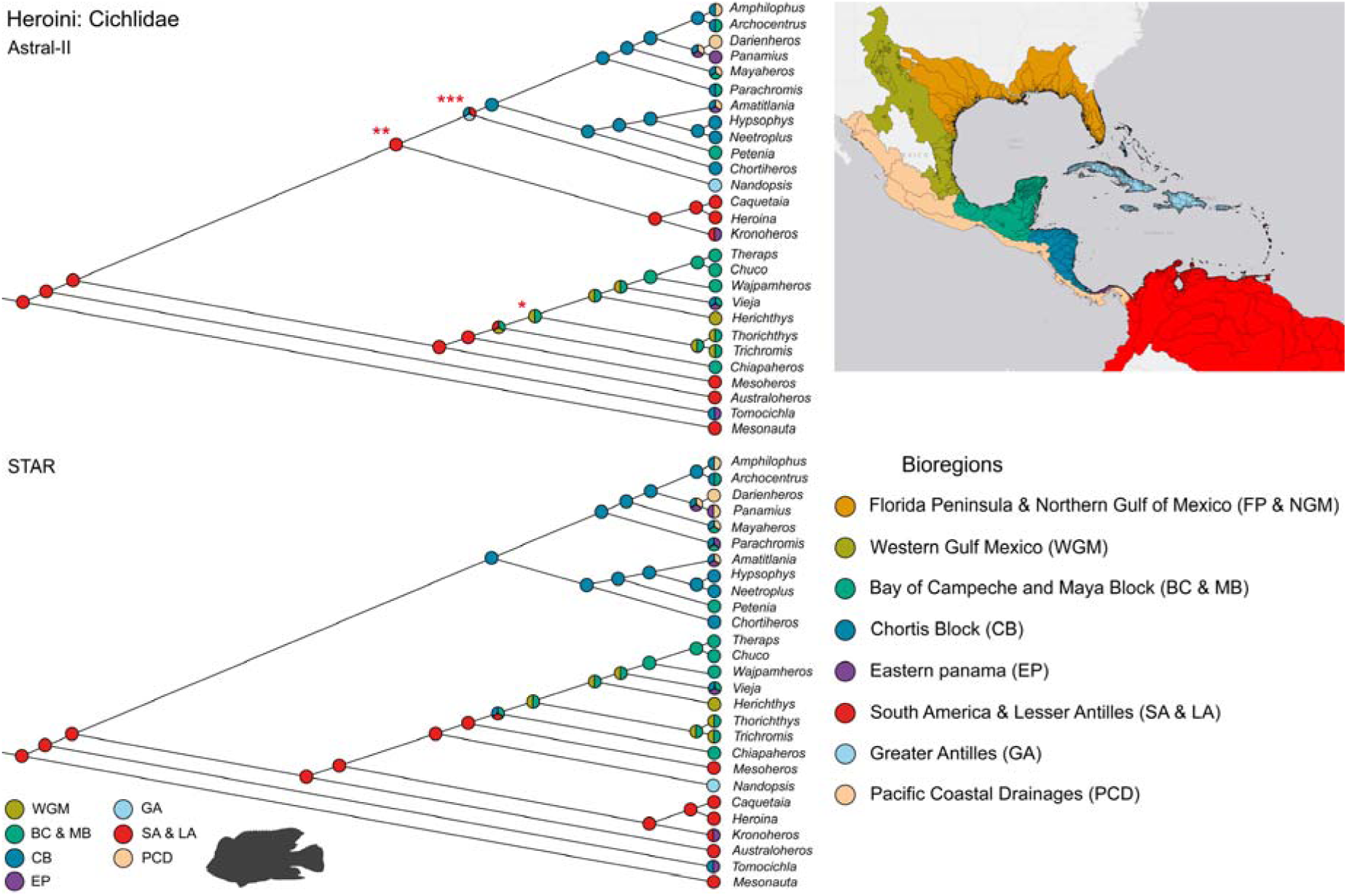
Final ancestral area analysis redrafted from Ilves et al. (2018) for the Heroini tribe, Cichlidae. Trees were obtained using ASTRAL-II (upper) and STAR (bottom) species trees methods with multi-locus bootstrapping. The position of Antillean *Nandopsis* changes radically depending on method. Asterisks denote weakly supported relationships.

**Fig. 8.**
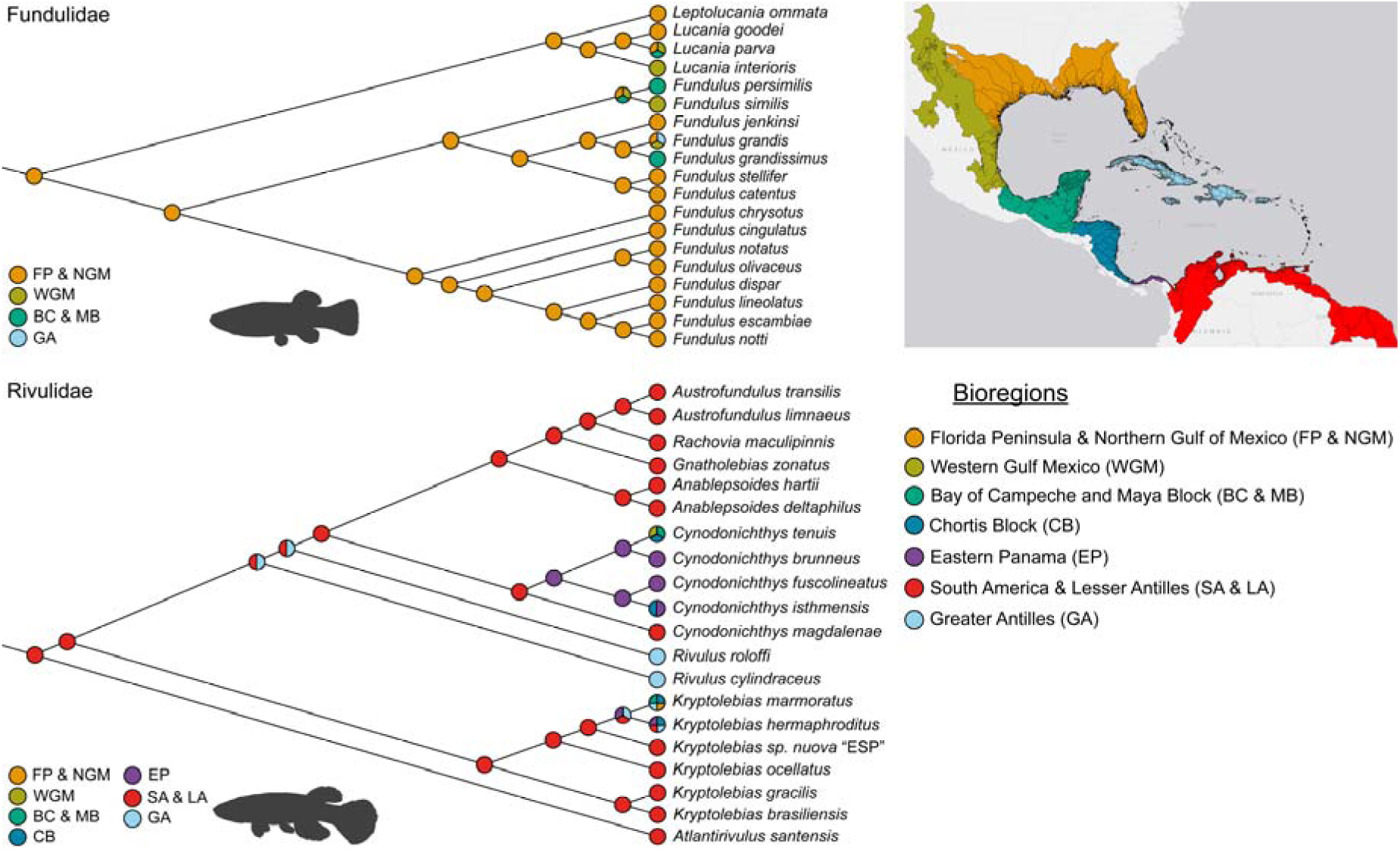
Final ancestral area state analysis redrafted from the phylogeny of Ghedotti & Davis (2017) for Fundulidae (upper) and our constructed phylogeny for Rivulidae (bottom).

**Fig. 9.**
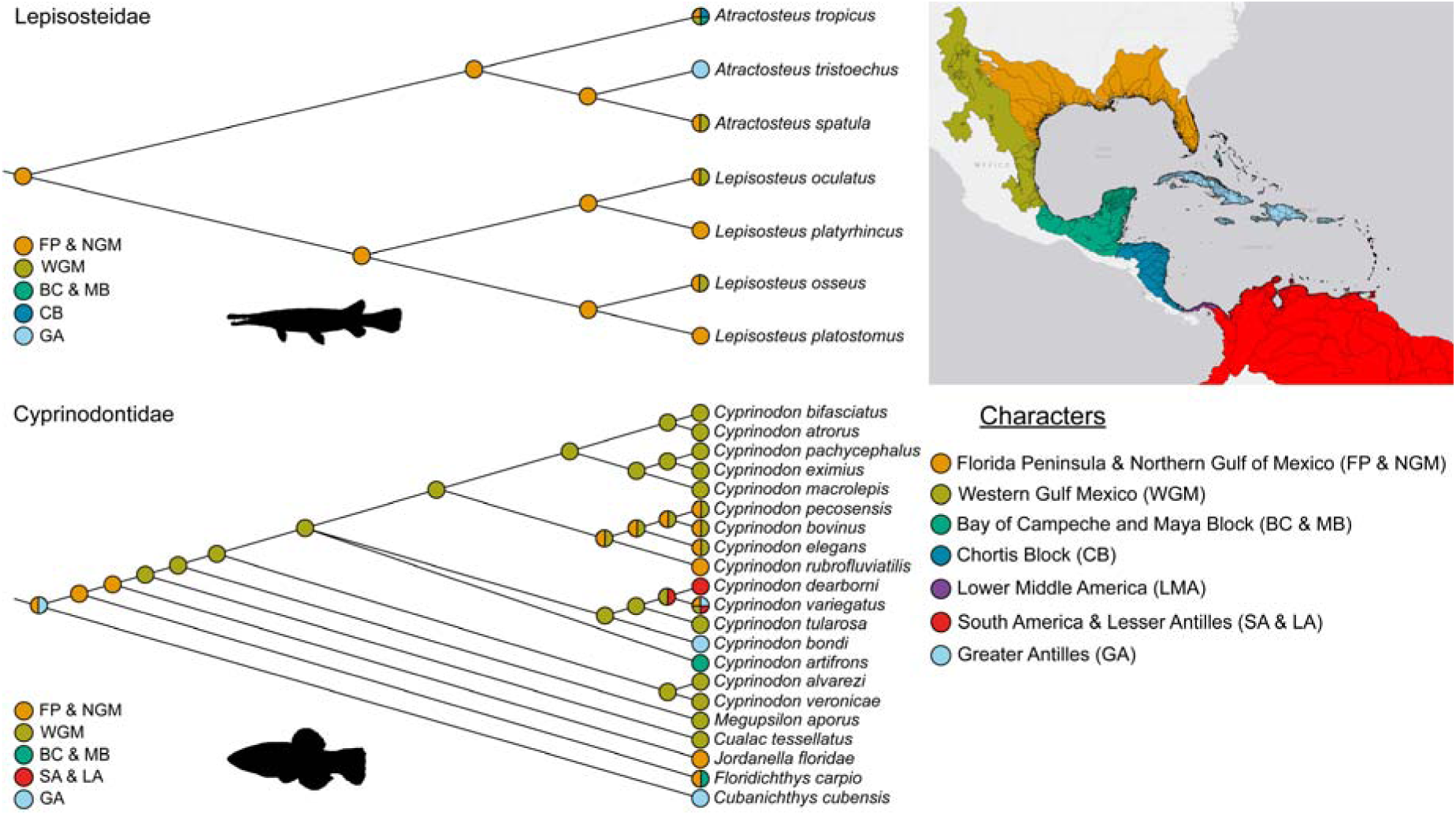
Final ancestral area state analysis based on the topology of Wright et al. (2012) for Lepisosteidae (upper) and our constructed phylogeny for Cyprinodontidae (bottom).

Because readers may conclude that we are biased defenders of long-distance-dispersal, we feel the need to state that this untrue. In this study, immigrations by cichlids and *G. melapleura-G. wrayi-G. hispaniolae* could be classified as a sort of vicariance if progenitors had contiguous distributions along the northern Nicaragua Rise when the Caribbean-Loop current was absent. In each case, establishment of the current across the Nicaragua Rise may have subdivided populations between Central America and the Greater Antilles. We view dispersal and vicariance as complementary. Vicariance occurs as barriers form, cross-barrier dispersal is required after barriers are formed. The longer a barrier exists, the more time there is for cross-barrier dispersal to happen and for any original, vicariant assemblage to disintegrate via extinction or shifting distributions. The antiquity of dispersal barriers surrounding the Greater Antilles creates a scenario potentially skewed toward long-distance dispersal, even if there were Late Cretaceous and Early Oligocene land connections.

Several authors have argued that rafts of vegetation discharged from large tropical rivers can facilitate overseas dispersal (Hedges, 2006; O’Dea et al., 2016; Ali, Fritz, & Vargas-Ramirez, 2021). Large rafts occur episodically and their dispersal may require strong winds or currents (Thiel & Haye, 2006). Although rare, favourable conditions appear to sometimes align along potential dispersal routes (Measey et al., 2007; Balaguera-Reina et al., 2021). Relevant examples in the Caribbean include freshwater turtles, snakes, and crocodilians (Charles, 2013; Brown et al., 2021). There appears to have been little (if any) observation of freshwater fishes associated with rafts in the ocean (Thiel & Gutow, 2005), yet within freshwaters, diverse fish assemblages disperse with rafts (Schiesari et al., 2003; Bulla et al., 2011; Dias et al., 2011). It seems likely that rafts discharged to the sea sometimes retain freshwater fishes or their propagules. If so, species that tolerate saltwater and harsh conditions should be most likely to persist. If a raft makes landfall, associated fishes could disperse along shore rather than being beached with the raft. If present when a raft is discovered, small-bodied fishes might be inconspicuous to observers or presumed native. Seagrass beds, salt marshes, macroalgal belts, and mangroves produce floating substrata that promote frequent rafting (Thiel & Haye, 2006). Freshwater fishes that occur broadly across the Greater Antilles and adjacent bioregions (*C. variegatus*, *F. grandis*, *G. rhizophorae*, *K. marmoratus*, *K. hermaphroditus*) occupy these habitats (Nordlie, 2006; Taylor, 2012; Lozano-Vilano & de la Maza-Benignos, 2017), which could help account for their broad distributions and documented gene flow across ocean gaps. This area is ripe for future research.

Rivas (1986) argued that a comprehensive explanation for biogeography of Antillean freshwater fishes should account for the depauperate freshwater-fish fauna on Puerto Rico. Furness, Reznick, & Avise (2016) delineated a cut-off in island size (4728 km^2^), below which extinction dominates community dynamics. Puerto Rico (8896 km^2^) is well above their threshold and is integral within the hypothesized GAARlandia landspan (Iturralde-Vinent & MacPhee, 1999; Iturralde-Vinent, 2006) and GrANoLA land mass (Philippon et al., 2020), implying these hypotheses are irrelevant for freshwater fishes. Studies to explain the lack of freshwater fishes on Puerto Rico could be informative for understanding freshwater-fish immigration. One possibility is that Puerto Rico is relatively remote from other bioregions except NSA and is isolated from that bioregion by the Caribbean Current, which flows westward, away from Puerto Rico. The one species inhabiting Puerto Rico, *K. hermaphroditus*, also inhabits the Lesser Antilles (Berbel-Filho et al., 2022), suggesting it dispersed along the Antillean Current from north-eastern South America, a pattern seen in juvenile sea turtles (Gaspar et al., 2022), but not other freshwater fishes.

## IV. Conclusions

1. Collectively, strong compositional disharmony, multi-way filtering of immigrants from diverse ancestral areas, and reciprocal illumination between phylogenetic and geologic evidence support dispersal (not vicariance) as the mode of immigration for freshwater fishes of the Greater Antilles. Although not all immigrations are well understood due to limited information, several align with conditions that could have facilitated oceanic dispersal and only one (*Girardinus-Quintana*) tentatively aligns with a landspan (GAARlandia), but the ancestral area of this group is uncertain. Of undated immigrations, only those for Cuban and Hispaniolan *Rivulus* are likely to bear on landspan hypotheses, because *Rivulus* is the only undated group with South America as its ancestral area. Thus, although we expect future research to revise these findings, it seems unlikely that a distinct pattern of landspan dispersal will emerge.
2. The GAARlandia hypothesis was proposed for terrestrial mammals (Iturralde-Vinent & MacPhee, 1999) and relies on evidence of Late Eocene Aves Ridge uplift (MacPhee & Iturralde-Vinent, 2005). Without evidence for freshwater connections along the proposed landspan, this hypothesis has limited implications for freshwater fishes and, with new evidence of Aves Ridge subsidence (instead of uplift) in the Late Eocene (Garrocq et al., 2021), the overall validity of the hypothesis is uncertain.
3. Physical and biological phenomena are integrative and interactive. As an example, we propose oceanic dispersal by freshwater fishes to the Greater Antilles relied on physical processes such as those producing rafts of vegetation (storms, floods) and delivering rafts to islands (ocean currents) along with biological factors (salinity tolerance, viviparity) to establish island populations. This is not a rejection of geology as important within biogeography, but a recognition that physical processes are more diverse than just plate tectonics and living things are not entirely passive (they struggle to exist).
4. Despite a conclusion that dispersal is the mostly likely mode of immigration for Antillean freshwater fishes, this does not imply oceanic dispersal is common or even likely. All evidence is consistent with its rarity over millions of years, even for better-adapted fishes under relatively favourable conditions. Nonetheless, hypotheses of dispersal need not be more ambiguous than hypotheses of vicariance. Reasoned predictions can be made regarding which taxa are most likely to disperse successfully, based on biological evidence. Physical conditions likely to favour dispersal can also be predicted and investigated. Studies in phylogeography can be particularly useful in this regard, when paired with hypotheses of dispersal and vicariance that are based on physical evidence. Studies of dispersal patterns in widespread taxa (which are increasingly plausible given emerging technologies) could also be highly informative.

## V. Acknowledgements

This paper is part of the requirements of the first author for obtaining a Doctoral degree at the Doctorado en Ciencias en Biodiversidad y Conservación de Ecosistemas Tropicales, Universidad de Ciencias y Artes de Chiapas (UNICACH) and received fellowship No. 833085 from Consejo Nacional de Humanidades, Ciencias, y Tecnologías (CONAHCYT). This project was partially supported by CONAHCYT grant CB-747-268.

## VII. Supporting information

**Appendix S1.** Freshwater fish taxa per island, area, percentage of species richness

