## Appendix S1 for "Biogeography of Greater Antillean freshwater fishes, with a review of competing hypotheses": Appendix S1.docx

Appendix S1. Freshwater fish taxa per island, area, species richness per island and percentage of total species richness.

| Species | Cuba | Hispaniola | Bahamas | Jamaica | Puerto Rico | Cayman Islands |
| --- | --- | --- | --- | --- | --- | --- |
| *Atractosteus tristoechus* | 1 | 0 | 0 | 0 | 0 | 0 |
| *Cubanichthys cubensis* | 1 | 0 | 0 | 0 | 0 | 0 |
| *Cubanichthys pengelleyi* | 0 | 0 | 0 | 1 | 0 | 0 |
| *Cyprinodon bondi* | 0 | 1 | 0 | 0 | 0 | 0 |
| *Cyprinodon brontotheroides* | 0 | 0 | 1 | 0 | 0 | 0 |
| *Cyprinodon desquamator* | 0 | 0 | 1 | 0 | 0 | 0 |
| *Cyprinodon higuey* | 0 | 1 | 0 | 0 | 0 | 0 |
| *Cyprinodon laciniatus* | 0 | 0 | 1 | 0 | 0 | 0 |
| *Cyprinodon nichollsi* | 0 | 1 | 0 | 0 | 0 | 0 |
| *Cyprinodon riverendi* | 1 | 0 | 0 | 1 | 0 | 1 |
| *Cyprinodon variegatus* | 1 | 0 | 1 | 1 | 0 | 1 |
| *Fundulus grandis* | 1 | 0 | 0 | 0 | 0 | 0 |
| *Gambusia beebei* | 0 | 1 | 0 | 0 | 0 | 0 |
| *Gambusia bucheri* | 1 | 0 | 0 | 0 | 0 | 0 |
| *Gambusia dominicensis* | 0 | 1 | 0 | 0 | 0 | 0 |
| *Gambusia hispaniolae* | 0 | 1 | 0 | 0 | 0 | 0 |
| *Gambusia hubbsi* | 0 | 0 | 1 | 0 | 0 | 0 |
| *Gambusia manni* | 0 | 0 | 1 | 0 | 0 | 0 |
| *Gambusia melapleura* | 0 | 0 | 0 | 1 | 0 | 0 |
| *Gambusia monticola* | 1 | 0 | 0 | 0 | 0 | 0 |
| *Gambusia pseudopunctata* | 0 | 1 | 0 | 0 | 0 | 0 |
| *Gambusia punctata* | 1 | 0 | 0 | 0 | 0 | 0 |
| *Gambusia puncticulata* | 1 | 0 | 1 | 1 | 0 | 1 |
| *Gambusia rhizophorae* | 1 | 0 | 0 | 0 | 0 | 0 |
| *Gambusia wrayi* | 0 | 0 | 0 | 1 | 0 | 0 |
| *Gambusia xanthosoma* | 0 | 0 | 0 | 0 | 0 | 1 |
| *Girardinus creolus* | 1 | 0 | 0 | 0 | 0 | 0 |
| *Girardinus cubensis* | 1 | 0 | 0 | 0 | 0 | 0 |
| *Girardinus denticulatus* | 1 | 0 | 0 | 0 | 0 | 0 |
| *Girardinus falcatus* | 1 | 0 | 0 | 0 | 0 | 0 |
| *Girardinus metallicus* | 1 | 0 | 0 | 0 | 0 | 0 |
| *Girardinus microdactylus* | 1 | 0 | 0 | 0 | 0 | 0 |
| *Girardinus rivasi* | 0 | 1 | 0 | 0 | 0 | 0 |
| *Girardinus uninotatus* | 1 | 0 | 0 | 0 | 0 | 0 |
| *Kryptolebias marmoratus* | 1 | 1 | 1 | 1 | 1 | 1 |
| *Limia caymanensis* | 0 | 0 | 0 | 0 | 0 | 1 |
| *Limia dominicensis* | 0 | 1 | 0 | 0 | 0 | 0 |
| *Limia fuscomaculata* | 0 | 1 | 0 | 0 | 0 | 0 |
| *Limia garnieri* | 0 | 1 | 0 | 0 | 0 | 0 |
| *Limia grossidens* | 0 | 1 | 0 | 0 | 0 | 0 |
| *Limia hispaniolana* | 0 | 1 | 0 | 0 | 0 | 0 |
| *Limia immaculata* | 0 | 1 | 0 | 0 | 0 | 0 |
| *Limia melanogaster* | 0 | 1 | 0 | 1 | 0 | 0 |
| *Limia melanonotata* | 0 | 1 | 0 | 0 | 0 | 0 |
| *Limia miragoanensis* | 0 | 1 | 0 | 0 | 0 | 0 |
| *Limia nigrofasciata* | 0 | 1 | 0 | 0 | 0 | 0 |
| *Limia ornata* | 0 | 1 | 0 | 0 | 0 | 0 |
| *Limia pauciradiata* | 0 | 1 | 0 | 0 | 0 | 0 |
| *Limia perugiae* | 0 | 1 | 0 | 0 | 0 | 0 |
| *Limia rivasi* | 0 | 1 | 0 | 0 | 0 | 0 |
| *Limia sulphurophila* | 0 | 1 | 0 | 0 | 0 | 0 |
| *Limia tridens* | 0 | 1 | 0 | 0 | 0 | 0 |
| *Limia versicolor* | 0 | 1 | 0 | 0 | 0 | 0 |
| *Limia vittata* | 1 | 0 | 0 | 0 | 0 | 0 |
| *Limia yaguajali* | 0 | 1 | 0 | 0 | 0 | 0 |
| *Limia zonata* | 0 | 1 | 0 | 0 | 0 | 0 |
| *Nandopsis haitiensis* | 0 | 1 | 0 | 0 | 0 | 0 |
| *Nandopsis ramsdeni* | 1 | 0 | 0 | 0 | 0 | 0 |
| *Nandopsis tetracanthus* | 1 | 0 | 0 | 0 | 0 | 0 |
| *Poecilia caudofasciata* | 0 | 0 | 0 | 1 | 0 | 0 |
| *Poecilia dominicensis* | 0 | 1 | 0 | 0 | 0 | 0 |
| *Poecilia elegans* | 0 | 1 | 0 | 0 | 0 | 0 |
| *Poecilia nicholsi* | 0 | 1 | 0 | 0 | 0 | 0 |
| *Quintana atrizona* | 1 | 0 | 0 | 0 | 0 | 0 |
| *Rivulus berovidesi* | 1 | 0 | 0 | 0 | 0 | 0 |
| *Rivulus cylindraceus* | 1 | 0 | 0 | 0 | 0 | 0 |
| *Rivulus roloffi* | 0 | 1 | 0 | 0 | 0 | 0 |
| Area (Km^2^) | 110860 | 79192 | 13880 | 10991 | 9104 | 260 |
| Species Richness | 24 | 33 | 8 | 9 | 1 | 6 |
| % of Total Species | 35.82 | 49.254 | 11.94 | 13.433 | 1.492 | 8.955 |
